# Thermodynamic principles govern evolutionary tradeoffs by regulating allostery

**DOI:** 10.1101/2025.10.08.680613

**Authors:** Yusran A. Muthahari, Reza Aditama, Mary Providaki, Alexandra Tsirigotaki, Chara Sarafoglou, Ruixue Xu, Srinath Krishnamurthy, Rukman Hertadi, Michael Kokkinidis, Cang Hui, Paola Laurino, Charalambos Pozidis, Giorgos Gouridis

## Abstract

Allostery governs biological activities by signaling environmental cues at distal sites, however, the molecular basis for the orchestration and evolution of “the second secret of life” is evanescent. Seminal work sheds light embracing structure- and function-centric approaches, overlooking the widely accepted ensemble allosteric model based on a proteins’ free-energy landscape and the thermodynamic nature of allostery. Here, we unraveled allosteric regulation and its evolvability by examining energetic funnels of proteins harboring a highly evolvable ancient scaffold. We uncover intricate allosteric connectivities and their coordinated cross-talk to enable the statistical thermodynamic coupling. We decipher universal molecular determinants for the emergence of functions and environmental adaptability. Our integrative biophysical/statistical/evolutionary analysis ties the evolutionary forces *via* thermodynamic principles and decrypts how tradeoffs are settled at the molecular level.

**One-Sentence Summary:** We reveal how physical and evolutionary laws shaped the biophysical properties of proteins

## Main Text

Allostery denotes the long-range modulation of protein activities by effector molecules. In 1904, the influence of carbon dioxide on the interaction of oxygen to haemoglobin was termed the “Bohr Effect” (*1*). About 60 years later, Monod, Wyman and Changeux formulated the first model to describe allostery (*2*). Since then, numerous others have been proposed for allosteric signal propagation (*3-7*). Current ones, based on the statistical nature of the protein structure rely on the notion of allosteric ensembles (*8-10*). The Ensemble Allosteric Model (EAM) proposes allosteric signal propagation *via* large-scale structural changes (Tier-0 dynamics) or by local structural fluctuations and order-to-disorder transitions (Tier-1/2 dynamics) (*9*). Such multi-Tier dynamics can be explored by assessing the Free-Energy Landscape (FEL) of proteins (*4, 11*), leveraging multiple biophysical approaches (*12*). Signal propagation stems from an uneven distribution of protein stability/rigidity throughout its structure reallocated *via* a chain of cooperative interactions (*13*).

Seminal work started to address the means by which allostery advances throughout evolution. Allosteric regulation has been shown to progress gradually, *via* latent allosteric networks responsible to generate a pool of phenotypic diversity (*14, 15*), anticipated to stem from rugged FELs (*14*). Effector molecules have been suggested to harness such a latent network during evolution to direct it to specific phenotypic trajectories. These observations are in line with the proposal on the emergence of new functions from generalist ancestors (*16, 17*). The prior work (*14, 15*) relied primarily on protein structure and/or function. However, inadequate to unravel generic transmittance principles, as allostery is thermodynamic in nature involving changes in enthalpy and/or entropy, where communication across the protein is mediated *via* multi-Tier dynamics (*5, 9, 18*). Here, we determine all three allosteric components [i.e. structural, thermodynamic, FEL (*19*)] of an ancestral bilobed scaffold (*20*). We further uncover the allosteric network and its variations among homologues to settle evolutionary tradeoffs, resulting in functional diversification. Using an integrative biophysical, statistical and evolutionary analysis, we establish connections among sequence, structure, energy wells and phenotypic traits, and the intricate sequence-structure-allostery-FEL-function-evolution relationship (Fig. 1, A and B).

**Fig. 1.**
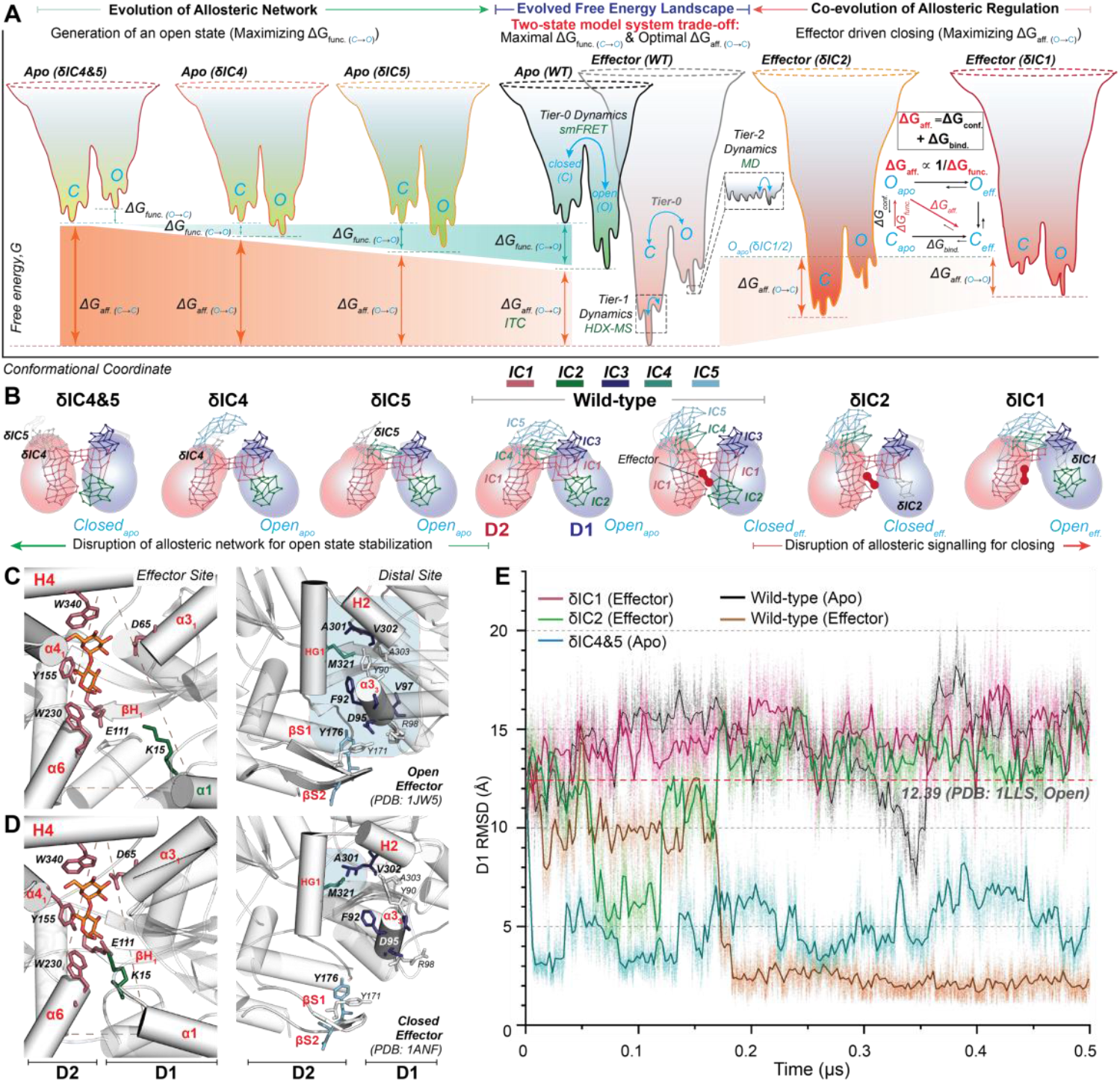
The allosteric network tailors the evolutionary tradeoff. (**A**) FEL of GCCPs and their thermodynamic cycle. Tier-0 “open” (O) and “closed” (C) states were determined by smFRET; Tier-1 and 2 by HDX-MS and MD respectively. Free energy differences between the lowest energy wells in the apo- and effector-FEL derived from ITC report on effector affinity (*ΔG*_*aff*. (O →C)_), whereas their free energy content via CD experiments inform on their thermodynamic stability. *ΔG*_*aff*_ can be decomposed in the conformational (*ΔG*_*conf*. (O →C)_) and binding (*ΔG*_*bind*. (C*apo* →C*eff*)_) free energy. *ΔG* between O and C in the apo funnel (*ΔG*_*func*. (C →O)_) determines GCCP functionality. *ΔG*_*aff*._ and *ΔG*_*func*_. are inversely proportional according to the thermodynamic cycle. Disruption of the allosteric network (i.e. δICs, as depicted) affect *ΔG*_*aff*._ and *ΔG*_*func*._ (**B**) Schematic representation of the ICs in the open or closed GCCP states. A and B contain key findings detailed in the main text. (**C, D**) Interactions between critical IC residues (color coded throughout as indicated in panel B) and/or the effector (orange sticks) are depicted for the open (C) and closed (D) MalE states. (**E**) All atom MD trajectories of the indicated derivatives. Open and the closed-effector bound state (PDB: 1ANF) were superimposed on D2, and the RMSD of D1 is visualized.

### The allosteric network of the bilobed scaffold

To uncover the Independent Components (ICs) of the allosteric sectors responsible for the allosteric event creating the two-state (open, closed) model system controlled by effector binding, we used the Statistical Coupling Analysis (SCA), introduced by Ernesto Freire and Rama Ranganathan (*21, 22*). We analyzed ∼4000 class-G Cherry Core Proteins (CCPs for the resemblance of the bilobed structure to a cherry) (*20*) (fig. S1A), with carbohydrates as effectors (GCCPs). The ICs were mapped onto the effector bound open (the two lobes or Domains D1&D2 are apart) or closed (D1&D2 come together akin a closed book) structures of the model GCCP protein (Maltose Binding Protein, MalE) used for biophysical characterization (Fig. 1, B to D and fig. S1). IC1 includes almost exclusively all D2 cleft residues responsible for effector docking, and a D1 residue that contacts the effector in the D2-docked (open) state (Fig. 1C and fig. S1, C to E). D65 is likely to be critical for allosteric signaling, being the only D1 residue contacting the effector in its docked state (fig. S1, C to E). IC2 embraces a D1 residue (K15) that interacts tightly with the effector solely in the closed state (Fig. S1, C to E). Three additional ICs (IC3-5), distal to the effector-cleft, are tightly interacting in the open state (Fig. 1, B and C and fig. S1F), whereas come apart in the closed (Fig. 1, B and D and fig. S1F), suggestive for open state stabilization. To verify if and how this IC network (fig. S1G) modulates allosteric transitions, we disrupted the corresponding ICs (table S1).

### Disruption of the allosteric network alters signal propagation

The IC3-5 interface was perturbed by simultaneously disrupting IC4 and IC5 (referred to as δIC4&5). All-atom Molecular Dynamic (MD) simulations (*23*) indicate that δIC4&5 bypasses the need of the effector for the allosteric transition (movie S3 *vs*. S2 and S1), as it switches intrinsically and readily (i.e., within 50 ns) from the open to the closed state (Fig. 1E). Contrary, in δIC1, allosteric propagation triggered by the effector is disrupted (Fig. 1E, movie S4 *vs*. S2). δIC2 is still proficient in conferring a two-state model system, yet the lifetime of the closed state is severely compromised (Fig. 1E, movie S5 *vs*. S2).

The capability of IC1-5 to govern the allosteric network was experimentally validated by single-molecule Förster Resonance Energy Transfer (smFRET) (Fig. 2, A to D) (*24*). D1&2 were labeled with donor and acceptor fluorophores to probe the two-state system (Fig. 1B) with single-molecule resolution (*20*). In line with our prior observations (*20, 25*), the unimodal distribution of low FRET efficiency values shifts towards higher values at saturating effector (maltose) concentrations (Fig. 2A). About equal amplitudes were observed for both distributions at effector concentrations proximate to the dissociation constant (*K*_d_) values (∼1,5 μM; Fig. 2A), with a ∼300 ms closed state lifetime after effector entrapment (*20, 25*). Remarkably, the two-state model system was abolished in δIC4&5 and δIC1, as unique closed or open states were observed respectively (Fig. 2, B and C and fig. S2, A and B). Additionally, δIC2 enabled acquisition of a closed state, however, requiring a ∼200-fold increase in the effector concentration (Fig. 2D), due to the decreased stability of the closed state (fig. S2, C and D). smFRET thus confirmed that IC1-5 are the master allosteric regulators. We next sought to enlighten the crosstalk between ICs underlying allosteric signaling.

**Fig. 2.**
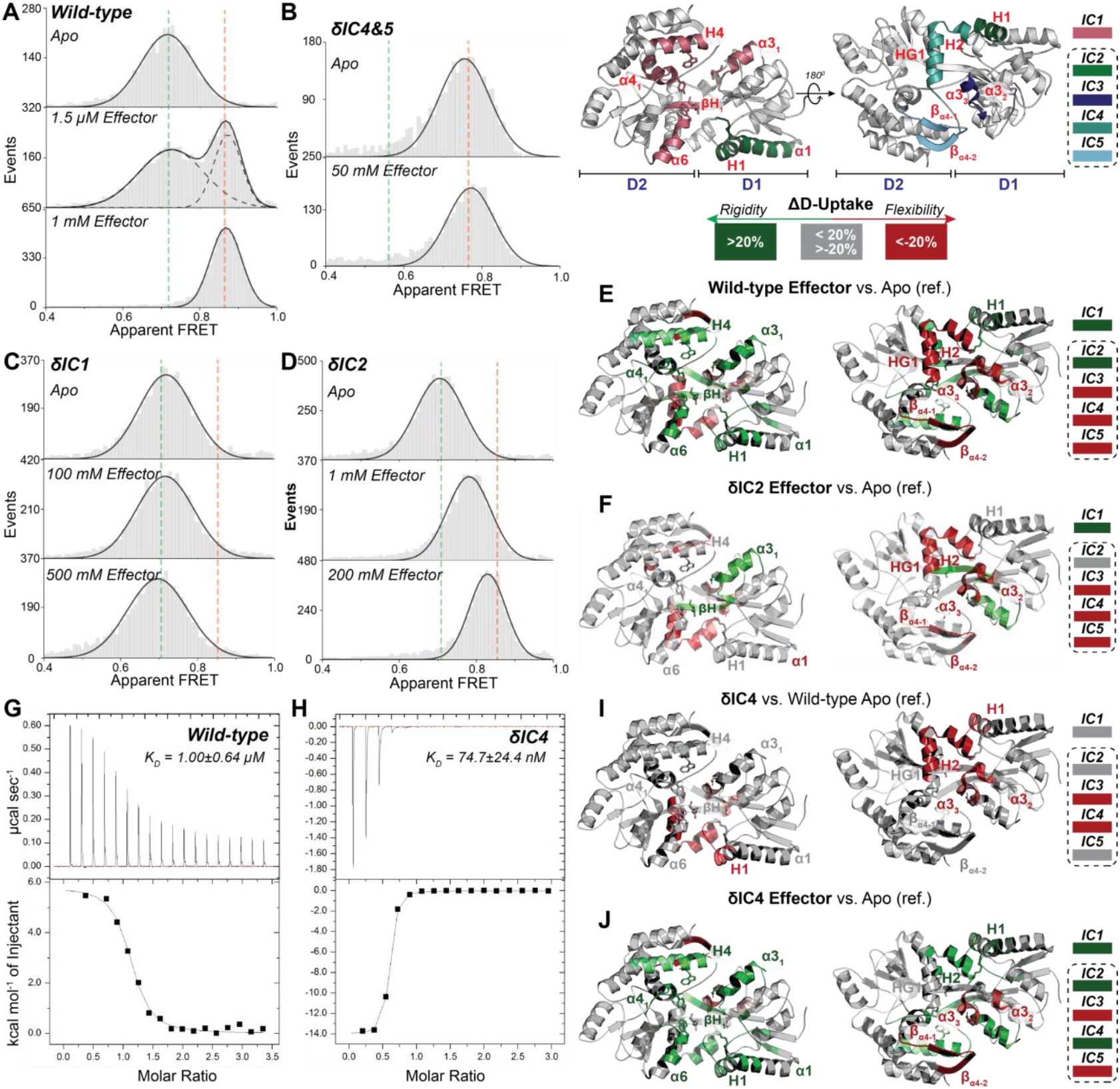
Disruption of the allosteric network impacts the structural and energetic landscape. (**A-D**) Tier-0 dynamics of wild-type and the derivatives probed by solution-based smFRET after titrating with different effector concentrations, as indicated. Green and orange vertical dashed lines denote the efficiency values for the open and the closed state of wild-type under identical microscope alignment conditions. (**E**-**G, I-J**) Structures showing the statistically significant differences in deuterium uptake caused by effector association or by allosteric network disruptions, as indicated (n=3). (**G-H**) Entropic and enthalpic binding isotherms after the calorimetric titration of the effector.

### The allosteric crosstalk enables the statistical thermodynamic coupling mediating signal propagation

The distribution of rigidity/flexibility within the bilobed structure was uncovered by Hydrogen Deuterium Exchange coupled to Mass Spectrometry (HDX-MS) (*26*). The ICs are characterized by islands with scattered stability (fig. S3 and fig. S4A). The structural fluctuations triggered by the allosteric event were decrypted by comparative HDX-MS and remarkably found to occur almost entirely on the ICs1-5 (Fig. 2E and Auxiliary files): The effector causes rigidification of IC1&2 (fig. S5, A and B) and a structural loosening of IC3-5 (fig. S5, C to E), following the allosteric transition from the open to the closed state. Flexible regions in the effector binding site and islands with low and high stability (i.e., α3_1_ and α4_1_ of IC1, respectively; fig. S3 and fig. S4A) have been shown to act as essential determinants for the allosteric signal propagation (*13*). Effector docking on IC1&2 (fig. S4B) modulates the structural dynamics of IC3-5 (fig. S4D *vs*. C and movie S2). To address this, we analyzed δIC1 or 2. The structural loosening of IC3-5 triggered by the effector is attained in δIC2 (Fig. 2F and fig. S4E), whereas completely impeded in δIC1 (fig. S4F), as no statistically significant difference in deuterium uptake occurred in all identified peptides (fig. S5F). Thus, IC1 is necessary and sufficient to drive allosterically the destabilization of the distal ICs, whereas IC2 controls the stability of the effector entrapped closed state (Fig. 1E and Fig. 2D and fig. S2, C and D) by modulating its effector-driven rigidification (fig. S5G). Furthermore, IC1 regulates the lifetime of the effector free closed state (fig. S4, G and H). Remarkably, δIC1 and δIC2 alter the intra- and inter-IC cross-talk (fig. S5, H and I) impeding a stable association of the effector (fig. S4I) and thus allosteric signaling. A pending question pertains to the impact of such rigidity-to-flexibility transitions on the energetics of effector association.

The thermodynamics of effector binding were followed by Isothermal Titration Calorimetry (ITC). The thermogram revealed an entropically-driven reaction, giving rise to a *K*_d_ of 1.00±0.64 μM (Fig. 2G). Such entropic contributions are likely to be caused by the structural loosening of the distal IC3-5, as revealed by HDX-MS (Fig. 2E). To evaluate this, we determined the energetics of effector binding in δIC1 or δIC2. Importantly, the entropic contributions vanished despite effector binding in δIC1, occurring however with significantly (∼10,000-fold) lower affinity (fig. S6, A and B). Consequently, δIC1 fails to signal the structural coupling responsible for loosening the distal ICs (fig. S5F *vs*. Fig. 2E and fig. S4F *vs*. fig. S4D and movie S4 *vs*. movie S2). This disruption prevents the allosteric transition from the open to the closed state, hence a unique open state is observed (Fig. 2C). In contrast, δIC2 that retains the ability to signal the detachment of the IC3-5 interface (fig. S4E and movie S5) and the concomitant structural loosening (Fig. 2F) leading to closing (Fig. 2D), preserves the entropic binding mode (fig. S6, A and C). We conclude that the entropic contributions, conferred *via* a structural loosening of the distal ICs3-5, are signaled by effector association on IC1 and its rigidification. Such a signaling cascade modulates the allosteric events to generate the two-state model system.

To uncover the molecular underpinnings of the effector-driven modulation of the conformational entropy by the distal ICs, we examined single and combined disruptions. δIC5 retained the entropically-driven mode of effector association (fig. S6, A and D). Remarkably, in δIC4, it switched to an enthalpically-driven reaction (Fig. 2H and fig. S6A). δIC4 caused a structural loosening of the IC3&4 peptides intrinsically (Fig. 2I and fig. S5, D and E), thus the entropic contribution fades in effector binding. Furthermore, the entropic contribution was unfavorable in δIC4 (fig. S6A), seemingly stemming from an effector driven rigidification of H2 in IC4 detected in HDX-MS (Fig. 2J and fig. S5J). Disruption of IC3-5, although distal from the effector binding cleft, affects the effector-driven enthalpy-entropy compensation and, by that, binding affinities (fig. S6A). In δIC4, although the entropic contribution became unfavorable for effector association, the affinity increased because of an enthalpic compensation. To understands such effects, we next sought to comprehend the means by which the ICs impact the FEL in two distinct chemical environments defined by the presence (saturating concentrations) or absence of the effector.

The thermodynamic stability of the open and closed states in the two FELs was assessed by Circular Dichroism (CD) via the extrapolation of melting temperatures (T_*m*_) (fig. S7). The free energies of the wild type structural states and of those having the ICs disrupted are represented in Figure 3A, conjointly with the Δ*G* between the lowest energy wells of the two FELs, derived from ITC measurements (fig. S6). Disruption of the proximal ICs (1&2), impacts the energy levels of the states in both FELs. In contrast, disruption of distal ICs (3-5) controls the levels of the states in the apoprotein-FEL. The Gibbs free energy levels dictate the effector affinity (fig. S6A and Fig. 1A) explaining the thermodynamics of binding (Fig. 3A). Specifically, in δIC4&5 the effector affinity increased by 3 orders of magnitude, despite the fact that the entropic contribution vanished and the reaction even became entropically unfavorable (fig. S6, A and E). In δIC4&5, the energetic level of the open state in the apoprotein FEL rise significantly, and by that the closed state became the predominant one (Fig. 2B). Since the energetic level of the closed state in the effector-FEL remained unaltered, the affinity increased considerably. Consequently, δIC4&5 caused a structural loosening intrinsically of the distal ICs [alike in δIC4 in Fig. 2I (*20*); and fig. S4J] and thus lacked the favorable entropic contributions.

**Fig. 3.**
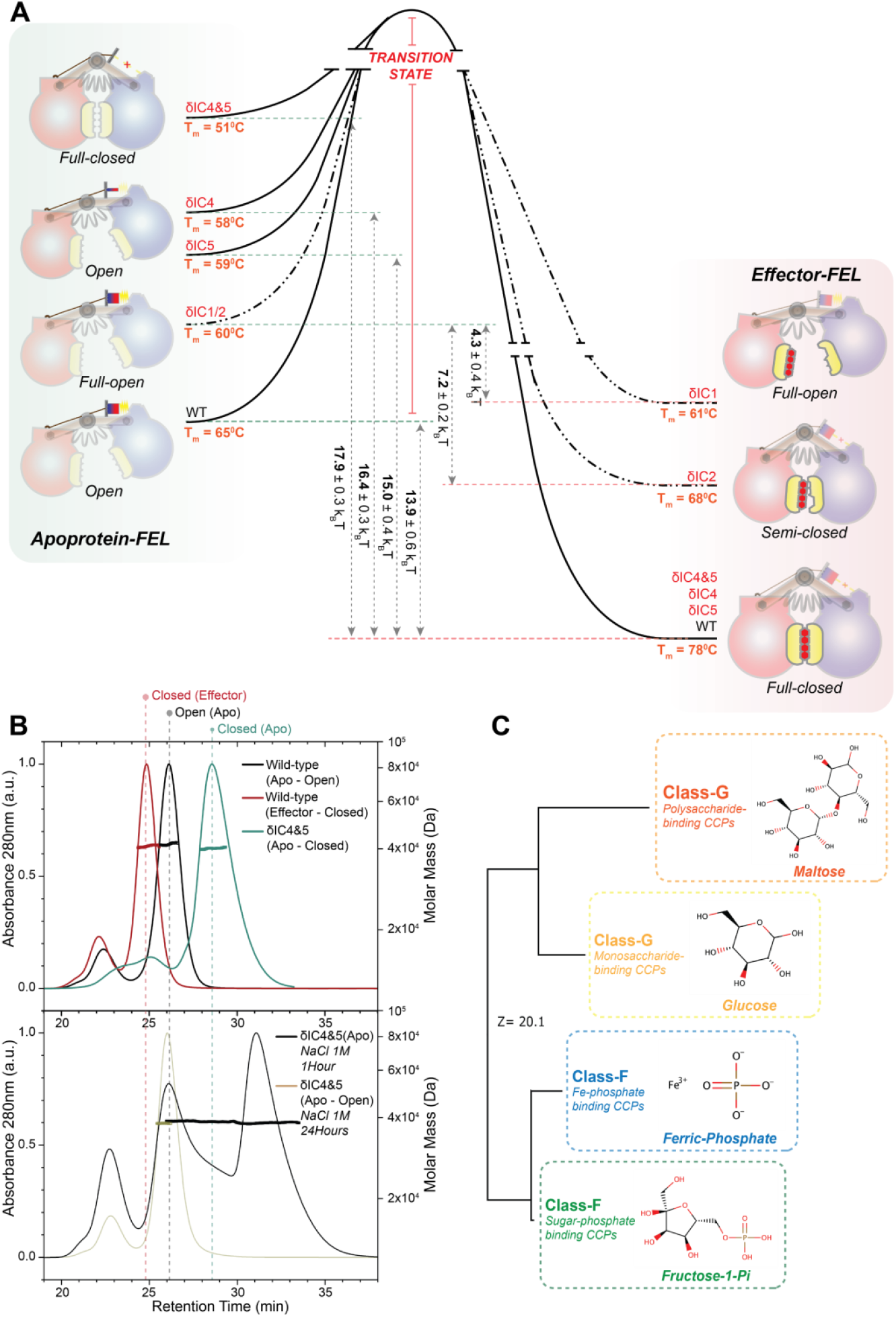
Thermodynamics govern the environmental and allosteric adaptation. (**A**) Schematic representation of the apoprotein (left) and effector (right) funnel wells together with the ITC and CD results. (**B**) Chromatograms coupled to MALS measurements of the indicated derivatives at physiological (top) and high (bottom) salt concentrations. (**C**) Structure-based phylogenetic tree with indicative class F & G CCP effectors.

### Perturbations on the allosteric network reflected in the thermodynamic parameters expose the “hidden” conformations that endorse protein evolvability

The thermodynamic analyses (Fig. 3A) implies the presence of multiple wildtype- and IC-disrupted energetic states, not just the two (open-closed) structural states deducted from smFRET (Fig. 2, A to D and fig. S2). All-atom MD trajectories corroborate the presence of multiple structural states, stabilized by interactions between the proximal ICs and/or the effector (fig. S4, K and L). The quantity of these interactions dictates the degree of closing (fig. S4L). The number of wildtype structural states in MalE is debatable, depending on the experimental tool used to probe them (*25, 27, 28*). To verify their occurrence, we opted to physically separate these states. Evaluation of the high-resolution structural information revealed two hydrophobic patches differentially exposed in open and closed states: one composed of IC1 residues and the second formed at the IC3-4 interface (fig. S8). MD revealed that the IC1 patch becomes progressively buried (fig. S4B), while the IC3-4 exposed (fig. S4D) during the allosteric transition to the closed state, controlled by the effector (movie S2). Concomitantly, IC1 is rigidified (fig. S5, A and B), whereas IC3-4 loosened (Fig. S5, C to E), as shown by HDX-MS (Fig. 2E). The separation relied on Size-Exclusion Chromatography (SEC) on Sepharose 75, which is known to separate species based on their degree of hydrophobicity (Fig. 3B and fig. S8) (*29, 30*). To verify that the separation depends uniquely on hydrophobicity, we coupled SEC to Multi-Angle Light Scattering (MALS) and performed a series of control experiments at different conditions that exaggerate (e.g., salt), mask (e.g., glycerol) or neglect (e.g., use of Superose 12) hydrophobicity (fig. S8). IC disruptions and thermodynamic conditions unmasked by separating a number of structural states (Fig. 3B and fig. S8C). As a case in point, the δIC4&5 state in the apoprotein-FEL has the same closed structure as the wildtype state in the effector-FEL (Fig. 3A), as revealed by smFRET (Fig. 2A-1mM-Effector *vs*. Fig. 2B-Apo). Using SEC-MALS, we fractionated the two closed (apo *vs*. effector-bound) states (Fig. 3B and fig. S8A), thus rationalizing their energetic difference (Fig. 3A). δIC4&5 has the IC3-4 hydrophobic patch intrinsically exposed (fig. S4J) and eluting at later time points compared to all other wildtype and disrupted states of the apoprotein-FEL (fig. S8D). Incubation with high salt concentration (1M NaCl) to maximize the hydrophobic effect, showcased a retarded elution (Fig. 3B, lower panel). However, a prolonged salt incubation (∼24h) was required for the system to reach equilibria (Fig. 3B and fig. S8C). At equilibria, the system has been shoved to hinder its hydrophobicity by obtaining the open state (Fig. 3B and fig. S8C). Thus, perturbations in the internal components (i.e., δIC4&5) and the chemical environment (i.e., salt) brought the system out of equilibria and delayed transition kinetics from the closed to the open state. This allowed us to expose a number (>5) of “hidden” structural states (fig. S8C, lower panel). Such “hidden” conformations (*16, 31, 32*) underlie the evolution of this bilobed scaffold, from a one-state (closed) to a two-state (open, closed) model system (*20*) (Fig. 1A). We have previously exposed these “hidden” structural states by exploiting promiscuous effectors [maltodextrins ranging from two to several glucosyl units in addition to maltose-(*25*)]. As shown here, the equilibria between the (hidden) structural states are controlled and can be biased by perturbing the ICs (fig. S4K).

### Alterations and expansions of allosteric networks settle evolutionary tradeoffs

The proximal IC1&2 that regulate effector binding (Fig. 1 and fig. S1 and fig. S6A), allosteric signal transmission (Fig. 2 and fig. S5) and concomitantly the FEL (Fig. 3A) are encompassed within the ancestral bilobed structure [fig. S1A and fig. S9, A and B and (*20*)]. We previously revealed that such a structure possesses a unique closed state displaying an effector capturing function (*20*) (i.e. ancestral trait). Consistently, the key IC1&2 residues (D65 and K15 in MalE) have coevolved to bind the different maltodextrin effectors in extant GCCPs (fig. S9, C and D). The distal IC3-5 have emerged as modular elements during evolution (fig. S1A and fig. S9A), diversifying the specificity and function of proteins harboring the ancestral bilobed structure (*20*). In line with our findings, mutations in distal ICs have been documented to affect allosterically the dynamics of effector clefts, conferring conditional neutrality in the PDZ domain architectures (*33, 34*) that are unrelated to the bilobed scaffold. Raman *et al*. proposed that neutrality acts as the source of functional diversification, paving the way for evolutionary adaptation (*33*). In the bilobed proteins, the diversification is accomplished by the “generation” of distinct open states (*20*). IC3-5 are specific to the class-G transport- and chemotaxis-related CCPs and generate an open state adapted to accommodate maltodextrin effectors (*20*), by controlling the apoprotein-FEL (Fig. 3A). A class-G subcategory entails short maltodextrin versions, i.e., monosaccharides, thus restricted effector-proximal IC1&2 interactions (Fig. 3C, see detailed explanation on the effector size; fig. S8) and consequently a diminished contribution from *ΔG*_*bind*_. (Fig. 1A). This increases the energetic level of the effector-FEL closed state. To maintain a physiologically relevant effector affinity (Fig. 1A), a corresponding increase in the energetic level of the open state in the apoprotein-FEL is foreseen. This indeed has been achieved during evolution by the deletion/insertion of the IC5 module (fig. S9, A and B). IC5 has been shown to stabilize the open state in the apoprotein-FEL (Fig. 3A), thereby increasing its energetic difference from the closed one. In line with these observations, class-F proteins, which are the most closely related to the class-G ones [Fig. 3C and fig. S9C and (*20*)], lack completely the extended class G ICs (4&5), while possessing a single restricted IC (ICX, fig. S9A). Class-F CCPs are typically associated with binding small-sized effectors as iron phosphates/carbonates. However, a class-F subcategory with bulky effectors (sugar-phosphates, spermidine/putrescine) or deriving from thermophiles, has detained IC4 (fig. S9). These findings are concordant to the antagonistic (i.e. beyond capturing) extant function of these CCPs, i.e., to mediate solute transport and chemotaxis through a firmly controlled two-state model system (extant trait) regulated by the effectors (*35*). Such a system lacks frequent intrinsic transitions to the closed state, which have been shown to severely compromise effector transport (*36*). Evidently, evolution has optimized the extant function by stabilizing the open state and by that extending the open-closed free-energy difference (*ΔG*_*func*_.) in the apoprotein-FEL (Fig. 1A). The difference has been widened even further in high-energy environments, as required, to prevent intrinsic transitions. This is, however, not possible when the effector is small or low-abundant (small contribution from *ΔG*_*bind*_.), given the fact that *ΔG*_*func*_. is inversely proportional to the affinity (*ΔG*_*aff*_., Fig. 1A).

## Conclusions

We have revealed that key allosteric residues coevolve with the effectors and are further expanded by the addition of allosteric modules that modulate the FEL and, in turn, fine-tune the evolutionary tradeoff (*37*) between transport/chemoreception and effector capturing capacity (Fig. 1, A and B). We anticipate such mechanisms to be widespread, as the majority of structures arise from the addition of elements to a common/conserved structural core, which constitutes ∼50% of the structure (*38*), alike to the CCPs (fig. S1A and fig. S9). We have, therefore, directly linked two evolutionary forces: the genetic diversity at the molecular level (*39*), by detecting its impact on structural dynamics, and the organismal-level selection, by associating dynamics to selected protein functionalities, effector specificities and environmental conditions emerged in related evolutionary pathways. Similarly, the molecular etiology of diseases can be derived via the linkage between disease symptoms at the cellular and allosteric action at the molecular level (*40*). As determining all allosteric components is fundamentally important for protein and drug design (*8, 41*), our findings need to be considered in the broader fields of molecular medicine and pharmacology besides protein engineering, biophysics and evolutionary biology.

## Acknowledgments

We thank S. Weiss and T. Cordes for providing software for data acquisition and analysis of ALEX data; A. Dömling and T. Economou for accessibility of infrastructure.

## Author contributions

Conceptualization: YAM, GG

Investigation: YAM, RA, MP, AT, CS, RX, SK, CP, GG

Formal Analysis: YAM, RA, MP, AT, CS, RX, RH, MK, CP, GG

Visualization: YAM, RA, AT, CS, CP, GG Supervision: RH, MK, CP, GG

Writing-original draft: YAM, GG

Writing-review & editing: YAM, RA, AT, CS, RH, MK, CH, PL CP, GG

## Data and materials availability

All data are available in the main text or the supplementary materials. Data analysis was accomplished by widely available published procedures and freely available software or servers as indicated in the main text or the supplementary materials

## Supplementary Materials

**PDF file that includes:**

Materials and Methods

Figures S1 to S9 including discussion and explanation

Tables S1 and S2

Caption for Movies S1 to S5

Caption for Auxiliary files

References (42–74)

Movies S1 to S5

Auxiliary files

