## Supplementary File for "Thermodynamic principles govern evolutionary tradeoffs by regulating allostery"

**This PDF file includes:**

Materials and Methods

Figures S1 to S9 including supplementary text for detailed explanation in each

Tables S1 and S2

Materials and Methods.

Statistical Coupling Analysis

Class-G CCPs structures were analyzed using the DALI server (*1*) as previously described (*2*). Using the PDB search options of the DALI server, we retrieved all the available high-similarity structures and evaluated them using the ECOD server (*3*). We manually curated proteins composed of the F-group SBP_bac_1_N_1 and SBP_bac_1_C domains to (a) select only the closely evolutionary-related proteins and (b) avoid excessive gaps in the Multiple Sequence Alignment (MSA). The PDB IDs of the selected structures are: 1OMP, 3JYR, 6L0Z, 2GBH, 6DTT, 6DTU, 2FNC, 6DTS, and 2GHA.

We generated the structure-based MSA by inputting the aforementioned structures to the DALI Server using the all-against-all feature. We expanded the MSA by inputting each of the selected structures to the protein-BLAST server (*4, 5*) with the default parameters (max target sequences: 5000). To prevent the excessive gaps in the MSA; we retrieved all protein sequences with similar length to the queries (±5 residues). All protein sequences were combined, the duplicates removed by SeqKit (*6*), and the remaining sequences were subsequently aligned to the structure-based MSA by ClustalX (*7*). The pySCA 6.1 (*8*) was performed to the final MSA by following the published methods (https://ranganathanlab.gitlab.io/pySCA/; accessed 20 January 2021) with 1OMP as the reference structure; however, we bypassed the MSA annotation process as such is not required in this study. The resulting ICs are mapped to different states of MalE structures to reveal the ICs’ structural connectivity (Fig. 1, B to D and fig. S1 and fig. S9).

Molecular Dynamics simulation

The crystal structure of the open MalE state is retrieved from Protein Data Bank with ID 1LLS, as described by Stockner (*9*). *In silico* mutations to disrupt IC1, IC3, IC4 were performed using PyMol. The structure of δIC5-containing derivatives was modeled by MODELLER (*10*) program server using the 1LLS structure as the template. The maltose used in the simulation is modified from the maltotetraitol structure retrieved from 1EZ9 (*9*). All simulations were carried out using AMBER 20. The proteins and maltose were represented using the FF14SB and GAFF forcefield, respectively (*11, 12*). The proteins were solvated in TIP3P water model with a distance of 1 nm to the edges of the box. Sodium ions were subsequently added to neutralize the system. The system was minimized and heated gradually to reach 305K while restraining the protein and maltose atoms with a restraint constant set to 10 kcal/mol Å^2^. The system was further equilibrated for 2 ns before releasing the positional restraint for the final 0.5 μs production run. During the production run, the temperature was kept constant at 305K with a Langevin thermostat. All the non-bonding interactions were confined within a 1 nm cutoff, where the electrostatic interactions were treated using the particle mesh Ewald (PME) method.

The trajectories were evaluated with the CPPTRAJ module of AMBER and subsequently analyzed with PyMol and VMD 1.9.1. To reveal the distinct states of MalE, all states were clustered using the KMeans algorithm after calculating the root mean square (rms) of D1 (residue number 1-105). The resulting clusters for each trajectory were manually analyzed and curated, revealing five different states of MalE.

Gene isolation, protein expression, and purification

MalE-His_6_ was expressed and purified as previously described (*2, 13*). *mal*E gene (UniProt: P0AEX9) was isolated from the genome of *Escherichia coli* K12. Primers introduced *Nde*I and *Hind*III restrictions sites, and the gene product was sub-cloned into the pET20b vector (Merck). MalE mutants were constructed using QuickChange mutagenesis.

BL21 DE3 cells (F– ompT gal dcm lon hsdSB(rB–mB–) λ(DE3 [lacI lacUV5-T7p07 ind1 sam7 nin5]) [malB+]K-12(λS)) was used to over-express MalE-His_6_. Cells harboring plasmids expressing the protein were grown in LB medium (37ºC; OD_600 nm_=0.8) and protein overexpression was induced by IPTG (0.5 mM; for 4 hours).

Harvested cells were diluted in 50 mM Tris-HCl, pH = 8; 1 M KCl, 10% glycerol; 10 mM Imidazole; 1 mM Dithiothreitol (DTT); 2 mM PMSF and subsequently lysed by the cell disruptor (30.000 psi; 2 rounds). After centrifugation at 50.000g for 30 min (4 ºC; Sorval), the soluble material was loaded on a Ni-NTA resin (Qiagen). Bound proteins were washed (50 mM Tris-HCl, pH = 8; 1 M KCl, 10% glycerol; 10 mM Imidazole; 1 mM DTT; and 50 mM Tris-HCl, pH=8; 50 mM KCl, 10% glycerol; 30 mM Imidazole; 1 mM DTT sequentially) and then eluted (50 mM Tris-HCl, pH=8; 50 mM KCl, 10% glycerol; 300 mM Imidazole; 1 mM DTT). Protein fractions were pooled (supplemented with 5 mM EDTA; 50 mM DTT), concentrated (Vivacell-Sartorious; Amicon-Millipore), dialyzed (50 mM Tris-HCl, pH=8; 50 mM KCl, 50% glycerol; 10 mM DTT), aliquoted and stored at -80ºC.

Protein labelling

MalE labelling was accomplished as described previously (*2, 13*). After performing structural analysis on the open and closed x-ray crystal structures of MalE (1OMP and 1ANF, respectively), residues number 36 and 352 were mutated into cysteines. MalE-His_6_ was immobilized on a Ni-NTA resin (Qiagen) in the presence of 1 mM DTT to maintain the cysteine's reduced state. The resin was incubated for 2-8 hours at 4°C with 50 mM Tris-HCl, pH=7.4, supplemented with 50nmol of Alexa Fluor 555 and Alexa 647 (ThermoFischer). The resin was subsequently washed to remove the majority of unbound fluorophores. The labelled protein was further analysed by size-exclusion chromatography (Superdex 200, GE Healthcare) to enrich the double-labelled fraction and remove potential aggregation material (*14*). For all proteins, labelling efficiency was higher than 80%.

Isothermal titration calorimetry (ITC)

Purified MalE was dialyzed against 1X PBS Buffer at 4 °C. ITC experiments of MalE were performed using MicroCal iTC200 (Malvern). Maltose solution was diluted in the dialysis buffer and was injected (2 μl) into the reaction cell containing 40 μM of MalE. Unless otherwise stated, all experiments were carried out at 280 K with a mixing rate of 750 rpm. Data (triplicates) from two independent protein purification procedures were analysed with a single-binding site equation provided by the MicroCal Analysis software (Malvern) and the standard error of the mean (SEM) was derived.

Multi-angle light scattering (MALS) and Quasi-elastic light scattering (QELS) experiments

Laser light scattering measurements were carried out online following size-exclusion chromatography (SEC) on a Superdex HR200 10/300 GL column and an HPLC system (LC10A-VP, Shimadzu) coupled to a quadruple detector scheme connected in series as follows: a photodiode-array detector (SPD- M10AVP; Shimadzu) for UV monitoring at 280 nm; an 18-angle MALLS detector (DAWN-EOS, Wyatt) using a K5-type cell and a laser wavelength of 690 nm; a QELS detector (WyattQELS; Wyatt) connected through an optical fiber to the MALS instrument through laser detector 13; and a refractive index detector (RID- 10A; Shimadzu). MALS and QELS data were collected, analysed and plotted using the Astra v.5.0 software (Wyatt). Proteins (2–100μM in 100μl injection volumes) in 50 mM Tris–HCl, pH 8.0, 50mM NaCl (or at the indicated NaCl and glycerol concentrations), were loaded onto the column and chromatographed at 22°C at a flow rate of 0.8ml min^-1^. Concentration at the peak fraction was typically 1/10 of the loaded concentration (data not shown). As for the experiments performed, samples obtained from two independent protein purification procedures produced chromatograms that overlaid perfectly with identical MALS output, one of the datasets is reported on Figure 3B and Supplementary Figure 8 for simplicity.

Circular Dichroism (CD) Measurements

Circular dichroism (CD) spectra (190–260 nm) were acquired using a J-810 CD spectropolarimeter (Jasco Inc., Easton, MD, USA) with a 1-mm path length quartz cuvette, at protein concentrations ranging from 0.1-0.3 mg/ml in 50mM Tris–HCl, pH 8.0, 50mM NaCl, supplemented with 15 mM β-mercaptoethanol. Thermal denaturation was analysed in the range of 10–90 °C in steps of 10 °C by monitoring the change of the signal at the typical α-helical minimum of 222 nm. Melting curves were obtained from the change of the CD signal at 222 nm in the above temperature range. Data (triplicates) together with the standard error of the mean (SEM) was determined by analysing samples deriving from two independent protein purification procedures.

The far-UV spectra were collected with 50 nm/min scanning speed, 1 min response time, and three accumulations. Thermal denaturation data were collected using a temperature increase of 80°C/h and a waiting time of 3 s for stabilization. The results were further analyzed by CalFitter webserver (*15*).

Solution-based smFRET and ALEX

ALEX experiments were carried out at 25-50 pM of double-labelled protein in the appropriate buffer (50 mM Tris-HCl, pH=7.4; 50 mM KCl) supplemented with additional reagents, as stated in the text. The experiments were performed using a home-built confocal microscope similar to the setup described before (*16*) with minor modifications as outlined below. Briefly, two laser-diodes (Coherent Obis) with emission wavelength of 532 and 637 nm were modulated in periods of 50 µs and used for confocal excitation. Alternation between both excitation wavelengths was achieved by direct modulation of the two lasers. The beam of both lasers was coupled into a single-mode fiber (PM-S405-XP, Thorlabs) and collimated (MB06, Q-Optics/Linos) before entering a water immersion objective (60X, NA 1.2, UPlanSAPO 60XO, Olympus). The excitation spot was focused 20 µm above the interface of glass and water solution. Typical average laser powers were 30 μW at 532 nm (~30 kW/cm^2^) and 15 μW at 637 nm (~15 kW/cm^2^). Excitation and emission light were separated by a dichroic beam splitter (zt532/642rpc, AHF Analysentechnik), mounted in an inverse microscope body (IX71, Olympus). Emitted light was focused onto a 50 µm pinhole and spectrally separated (640DCXR, AHF Analysentechnik) onto two APDs (τ-spad, <50 dark-counts/s, Excelitas Technologies) with appropriate spectral filtering (donor channel: HC582/75; acceptor channel: Edge Basic 647LP; both AHF Analysentechnik). Photon arrival times were registered by an NI-Card (PCI-6601, National Instruments). A dual-colour burst search (*17*) using parameters M = 15, T = 500 μs and L = 25 was applied to identify bursts. Additional thresholding was applied to remove spurious changes in fluorescence intensity and selected for intense single-molecule bursts (total photons per burst > 150 unless otherwise mentioned). Binning the detected bursts into a 2D apparent FRET/S histogram (61 x 61 bins unless otherwise mentioned) allowed the selection of the donor and acceptor labelled molecules (*18*). The selected apparent FRET histograms (61 x 1 bins unless otherwise mentioned) were fitted using a Gaussian function. Histograms were derived from data collected from two independent protein purification procedures that yielded more than 5000 single-molecules analysed per condition producing more than 150 Events (peak of the Gaussian function) per condition in order to fit unambiguously the Gaussian distributions.

Single-molecule bursts from donor-only labelled MalE proteins were obtained by selecting the donor-only subpopulation (S > 0.9) from the 2D apparent FRET/S histogram. The total photon count per burst were normalised by its respective duration to obtain the photon count rate.

Scanning confocal microscopy

The scanning experiments have been performed by using a home-built smFRET microscope that is equipped with an XYZ-piezo stage with a 100 × 100 × 20 µm range (P-517–3 CD with E-725.3CDA, Physik Instrumente), as described previously (*13, 19*). The HydraHarp 400 picosecond event timer and a module for time-correlated single photon counting (Picoquant) were used to register the detector signal. The data was acquired with 532 nm excitation (0.5 μW or ~125 W/cm2). After scanning images (10 × 10 µm) were recorded, the position of the labeled protein was retrieved and used to generate time traces. The protein surface immobilization has been performed by following the established protocol (*16*) without using any flow-cell arrangement to avoid maltodextrin contamination (*13*). The experiments have been conducted at room temperature using the same buffer used in the solution-based smFRET and ALEX with the addition of 1mM (±)−6-Hydroxy-2,5,7,8-tetramethylchromane-2-carboxylic acid (Trolox, Merck) (*20*). More than 500 single-molecules from two independent protein purification procedures per condition were collected with a total recording time of more than 10 min per condition.

Hydrogen/Deuterium exchange Mass Spectrometry (HDX-MS)

Isotope labeling: MalE and derivative δIC1/δIC2/δIC4 were thermally equilibrated for 30 min at 25^o^C [4 μl of 28.25 μM protein stock in 50 mM Tris-HCl pH 7.4, 50 mM KCl to ensure saturation and to thermally equilibrate the samples], in the presence or absence of 1 μl of 40 mM maltose (monohydrate grade 1, M-5885, Sigma Aldrich). For the apo state, 1μl of buffer was added instead. At 30 min of incubation, MalE was isotopically labeled with 93.75% v/v final D content at 25°C, by addition of 75 μl deuterated buffer [50 mM Tris-DCl pD 7.4, 50 mM KCl in D_2_O (D_2_O, 99.9% atom D; Euriso-top)] for 10, 100, 1000, 10000 and 100000 sec. pD refers to the corrected value for the isotope effect. The HDX reaction was quenched at the defined time intervals by instant acidification [pD 2.5; formic acid; (Ultra-pure) from Merck KGaA]. The pre-chilled quenching solution contained urea (Urea-d4; Sigma) to a final concentration of 1.6 M at quenching to increase the peptide coverage by mild denaturation. 50 pmoles of protein were injected for analysis.

Online proteolysis-LC-MS analysis: The quenched samples were injected into a nanoACQUITY UPLC System with HDX technology (Waters, UK), thermostated at the digestion and LC separation chambers at 20°C and 0.8 °C, respectively. Proteolytic digestion (Enzymate BEH pepsin column, Waters) and peptide trapping/desalting (ACQUITY UPLC R BEH C18 VanGuard pre-column; 130 Å, 1.7 μm, 2.1 x 5 mm; Waters) were performed with 0.23% formic acid in H_2_O (Solvent A) at 100 μl/min for 3 min, online with peptic peptide separation (ACQUITY UPLC R BEH C18 analytical column; 130 Å, 1.7 μm, 1 x 100 mm; Waters) at 40 μl/min using a 12 min linear gradient from 5% to 50% Solvent B [ACN (Optima LC/MS grade; Fischer Scientific), 0.23% formic acid]. The eluate was analyzed online on a Synapt G2 ESI-Q-TOF instrument (Waters, UK) with a MassLynX interface (version 4.1 SCN870; Waters) for data collection. The source/TOF conditions were set as: resolution mode, capillary voltage 3.0 kV, sampling cone voltage 20 V, extraction cone voltage 3.6 V, source temperature 80°C, desolvation gas flow 500 L/h at 150°C. The deuterated samples were analyzed in MS acquisition mode (300-2000 Da range), while for peptide identification non-deuterated samples (treated as above but in protiated buffers) were analyzed in MS^E^ acquisition mode over the m/z range 100-2,000 Da, using a collision energy ramp from 10 to 30 V. Leucine Enkephalin (2 ng/μl in 50% ACN, 0.1% formic acid; 5 μl/min) was co-infused in both acquisition modes for accurate mass measurements (reference mas: m/z 556.2771).

Data analysis: For peptide identification, MS^E^ data was processed on the ProteinLynx Global Server (PLGS v3.0.1, Waters, UK), using a user-defined database containing MalE, dIC1, dIC2, and dIC4 sequences under the following criteria: digestion enzyme, non-specific; false discovery rate, 4%, minimum fragment ion matches/peptide and /protein, 3 and 7; minimum peptide matches/protein, 1; low and elevated energy thresholds, 150 and 25 counts; intensity threshold: 500 counts, reference mass correction window, 0.25 Da at 556.2771 Da/e. The identified peptides from two independent MS^E^ raw files were further filtered on the DynamX software (version 3.0, Waters) for DynamX score> 7.5, maximum MH+ error of 5 ppm and minimum products/amino acid of 0.2. Only robustly identified peptides in both replicates were further processed, resulting in ~95% sequence coverage for MalE and disrupted derivatives. Deuterium uptake was determined using the DynamX software. For the comparison of the MalE and δIC4 at the mutated region, the peptide containing residues 321-337 was compared only at the level of D-uptake relative to the full deuteration of each state, after complete analysis of each protein with its individual controls, and not at the level of absolute D-uptake. For this, δIC4 was analyzed using the peptide list created from the δIC4 sequence, the δIC4 non-deuterated, and fully deuterated controls (Auxiliary files, Data S1B). Additionally, for the comparison of δIC4 at the apo state, statistical analysis was performed both on the data analyzed using the MalE and δIC4 sequence in order to include the mutated region (Auxiliary files, Data S1C-D). The comparisons between δIC1, δIC2 and MalE were performed similarly as above and reported in Auxiliary files, Data S1E. Analysis and interpretation of these states was carried out as described below.

Statistical analysis: All HDX reactions were performed in triplicates from two independent protein purification procedures unless otherwise indicated (Auxiliary files, Data S1). Statistical analysis of the significance of differences between the apo and effector-bound states, and between the MalE and δIC4 mutated sequence, was achieved using a modified approach of Bennett *et al*., (*21*), described in Tsirigotaki *et al*., (*22, 23*). Briefly, two-tail paired t-tests, comparing the mean uptake, as absolute deuterium uptake values, of the two states for each peptide, were performed using R language and the significance threshold was set to 99% confidence (1-p≥0.99). An additional threshold was set at ±4SD of average pooled standard deviations of both states. Finally, a third criterion was introduced to exclude false positives due to high SD outliers that are averaged out in the pooled SDs: The difference between the two states for each peptide must exceed twice the sum of SDs of the two states for the given peptide. Only differences that fulfill all three criteria were considered as statistically significant. Visualization of the statistical analysis in scatter plots was achieved by R language (Auxiliary files, Data S1D). For optimal realization of the extent of the ΔD-uptake regardless of the length of the identified peptides, the ΔD-uptake of each peptide was expressed relative to the experimentally determined full deuteration control (normalized ΔD-uptake). For higher robustness, only statistically significant differences that exceed the absolute value of 10% normalized ΔD-uptake are discussed. Detailed data sets, including smaller, statistically significant differences, are given in Auxiliary files, Data S1B for δIC4 apo and effector states; Data S1C for all comparisons between MalE and δIC4 at all states. Comparisons between MalE, δIC1 and δIC2 at all states were assessed based on two thresholds, 1) a 0.5 Da absolute difference in D-uptake, 2) a 10% normalized ΔD-uptake, calculated as described above. This two-threshold analysis ensured all differences satisfied a 99.5% confidence interval threshold. These data are tabulated and reported in Auxiliary files, Data S1E.

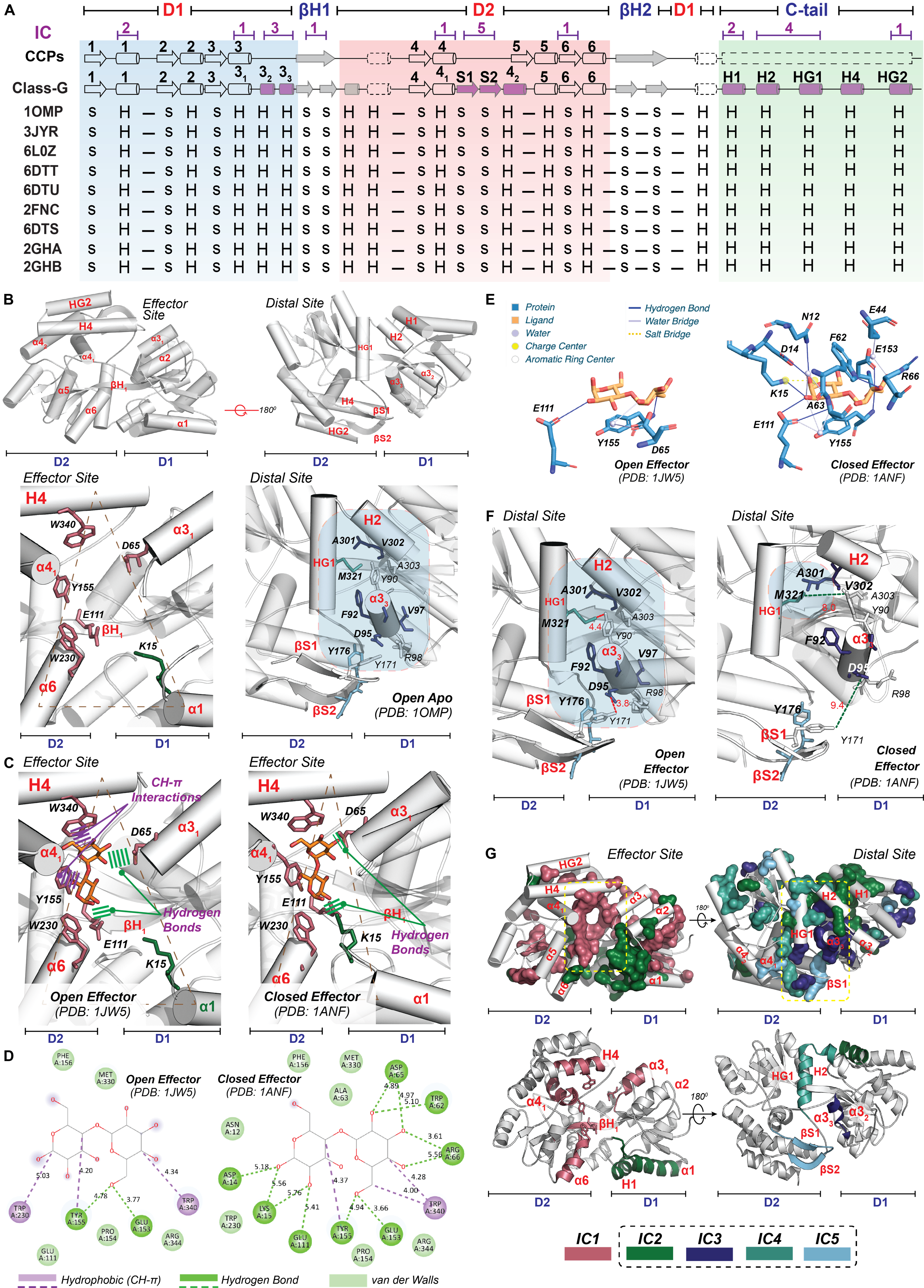

**Figure S1. Structural connectivity of GCCP ICs forming the allosteric network and critical IC residues regulating the two-state model system.** We choose MalE as our model GCCP structure for Independent Component (IC) visualization, structural and biophysical analysis; as it is the only GCCP for which complete high-resolution structural information is available for open- and closed-states in the presence and absence of the effector (i.e., maltose). The open, effector-free MalE structure (PDB: 1OMP) has been used as the reference structure for the Statistical Coupling Analysis (SCA) (*8, 24*). The resulting ICs were subsequently mapped onto the MalE crystal structures present in different states, allowing to guesstimate their function/role. (**A**) Secondary structure-based alignments to highlight the core (ancestral) and modular structural elements of GCCPs. The conserved core is shown in the top row as previously published (*2*) together with the ICs. GCCP structural elements colored in purple depict the modular ones added during evolution to the conserved core. (**B**) MalE structure (PDB:1OMP) showcasing key structural elements and residues (detailed in panels C-F) of the two domains (D1&2) of the bilobed structure. (**C**) The effector binding cleft (effector site) is formed by ICs belonging to two sectors. The first SCA sector includes a single IC [see (*25, 26*)], i.e., IC1. It consists of D65 (α3_1_), E111 (βH1), Y155 (α4_1_), W230 (α6), and W340 (H4). All cleft residues of IC1 are located mainly on D2, except for D65 situated on D1. (**D, E**) Maltose interactions with binding cleft residues represented in 2 dimensions (2D) and 3D, as evaluated by the Biovia Discovery Studio Visualizer (*27*) (D) and Protein Ligand Interaction Profiler (PLIP) webservers (*28*) (E), respectively; distances are shown in Angstrom. Based on the structural analysis of the effector-bound open state (PDB: 1JW5), several aromatic residues of IC1, i.e., W340 and Y155, enable initial effector docking on D2 (left side of panel C, D&E for 3D and 2D representation respectively). Indeed, aromatic residues are known to play a critical role in protein-carbohydrate recognition (*29*). The solvent-exposed aromatic ring of Y155 and W340 interacts with maltose via CH-π interactions (purple line panel C&D - left). Additional interactions are also provided by E111 (βH1) and E153 (α4_1_) to allow maltose-binding via hydrogen bonds (dark-green line in panel C&D; E153 is not shown in C for clarity). D1 residues responsible for effector association are derived from the structural analysis of the closed-liganded state (right C panel): D65 and K15 are the two residues that bind maltose (panel C **-** right). K15 is the second charged residue belonging to D1 that contacts the effector and is part of IC2 of the second sector. In the closed state, D65 forms two hydrogen bonds with the second ring of maltose (green line, panel C **-** right). Additionally, K15 (α1) stabilizes the closed state further by contacting both the effector and E111 (green line panel C **-** right). As D65 is the only D1 residue that ‘sense’ the presence of effector in the open state by contacting the effector (panel E), we hypothesized that D65 has a critical role in effector sensing triggering allosteric events. (**F**) The distal residues of the second SCA sector are tightly packed in the open state: The second SCA sector consists of 6 ICs, (i.e., IC2-7) and covers ~20% of the GCCPs. Interestingly, K15 of IC2 is the only residue in the effector cleft and is part of the ancestral α1 helix. The remaining ICs of the second sector are distributed throughout the GCCPs. IC3-5 are located distally from the cleft, whereas IC6&7 on the conserved core of the ancestral CC domains. As only IC3-5 undergo distance fluctuations in the open-closed states (dashed lines, distance in Angstrom), we postulated that only those regulate the two-state allosteric system (blue area in panel F **-** left *vs.* right). IC3 includes α3_3_ that expands the D1 structure at the backside of GCCPs and is positioned below the β-sheet hinges. Such a helix is located close to HG1 of IC4 and the βS1-2 hairpin motif of IC5. M321 of IC4 is buried by two aromatic residues from IC3, i.e., Y90 and F92. All the residues shown above are color-coded with respect to the ICs they belong to. The secondary structure elements numbering follows the topology of GCCPs (see also fig. S9). (**G**) All IC residues on the effector and distal sites are color-coded in the front and back views of the MalE structure as indicated (top panels). The critical residues of IC1-5 that emerged from the structural analysis detailed in panels B-F belong to specific secondary structure elements, as indicated in the bottom panels. Thus, we redefine ICs as the secondary structure elements indicated here to simplify the interpretation of the biophysical results throughout the manuscript.

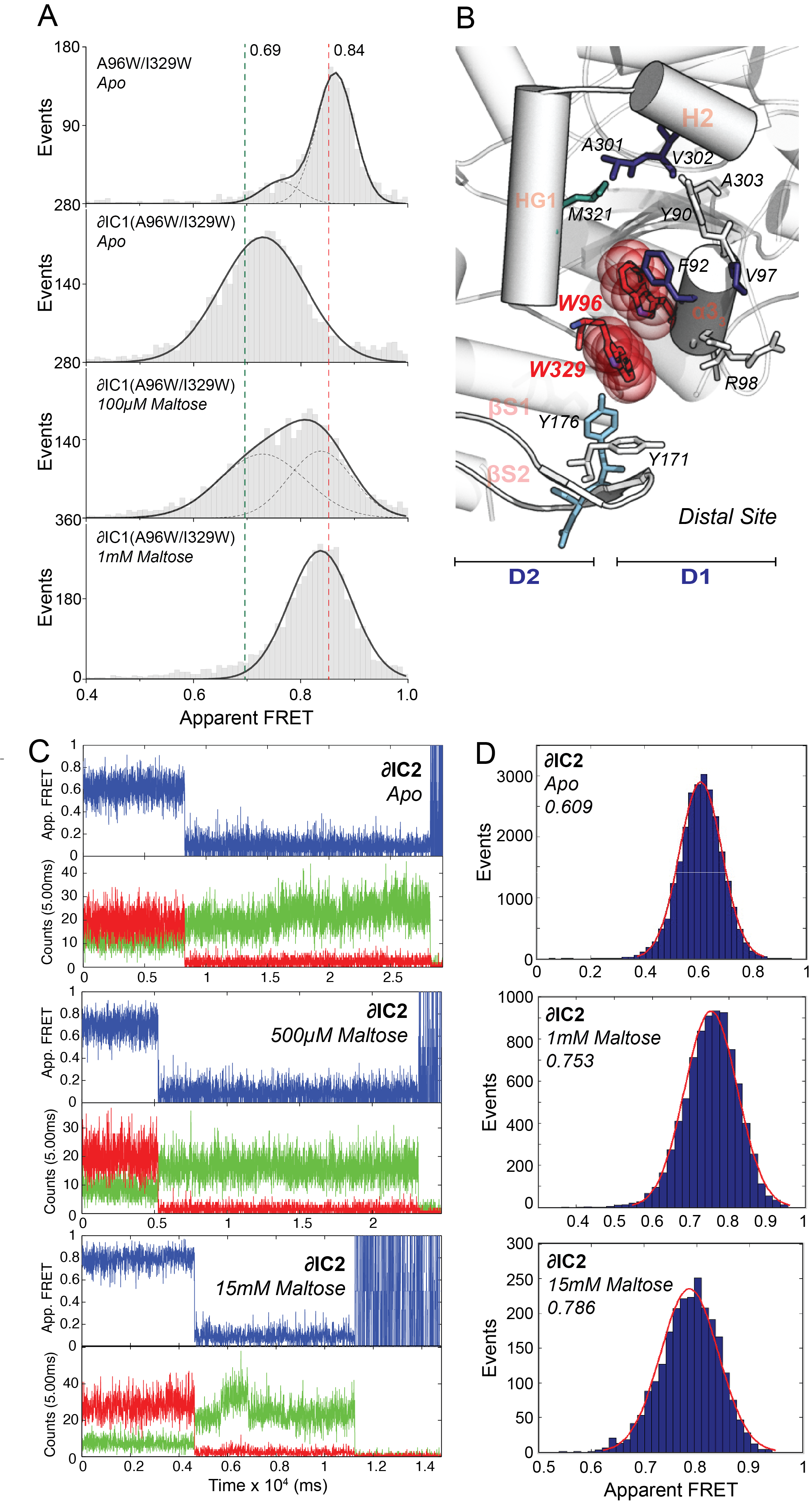

**Figure S2. smFRET reveals Tier-0 dynamics of GCCPs and the effect of IC-disruptions**. We performed single-molecule FRET to probe the tertiary dynamics of GCCPs. D1&2 of MalE were labelled with donor and acceptor fluorophores on T36 and S352 (*2, 13*). Notably, determination of anisotropy and ligand dissociation constants of labeled-MalE (*2, 13*) confirmed that FRET can act as a distance ruler to probe allosteric evens triggered by effector association. As detailed in the main text, 𝛿IC4&5 exhibits a unique closed state, irrespective of the presence of the effector (Fig. 2B). 𝛿IC1 possesses solely the open state without any observable structural change, even after effector addition (500mM maltose, Fig. 2C). (**A**) Solution-based FRET efficiency histograms of A96W/I329W and 𝛿IC1(A96W/I329W) titrated with different concentrations of the effector, as indicated. The green and red vertical dashed lines on the histograms denote the apparent FRET efficiency values of the open and the close state of MalE, respectively. (**B**) MalE(A96W/I329W) open state to visualize the steric clash between the two tryptophans. Alanine and isoleucine were replaced to tryptophans using Pymol. To analyze further the IC1 involvement in closing (intrinsic and effector driven), we made use of MalE (A96W/I329W) known to transit intrinsically to a closed state (*30*). Our findings provide a structural explanation of this phenotype: The bulky tryptophans in MalE(A96W/I329W) destabilize the IC3-5 interface (panel B, see also fig. S4G), alike in 𝛿IC4&5 (fig. S4J). MD showed that both MalE(A96W/I329W) and 𝛿IC4&5 transit readily to the closed state within ~50ns of the simulation (fig. S4H and fig. S4K, respectively). In line with published results (*30*), we observed the majority of the molecules present in a closed- effector free state (A, top panel). 𝛿IC1 abolishes acquisition of such a closed state. Remarkably, the addition of the effector, as indicated, triggers closing (A, bottom panels). This unexpected result (i.e., IC1 is necessary and sufficient for effector-driven closing as detailed in the main text) was disentangled by MD simulations. Despite the fact 𝛿IC1(A96W/I329W) cannot obtain a stable closed state (panel A) it maintains the ability to transit intrinsically to a short-lived (~50ns) closed state (fig. S4H). Taken together, these results indicate that IC1 is necessary and sufficient to trigger allosterically the closing event, and additionally critical to stabilize the closed state. **(C)** Indicative fluorescence trajectories of 𝛿IC2 surface tethered molecules at different effector concentrations as indicated; donor (green) and acceptor (red) photon counts are binned with 5ms. The top panel shows calculated apparent FRET efficiency (blue). Saturating effector concentrations trigger the transition of 𝛿IC2 to an almost fully closed state (see also Fig. 2D), however at intermediate concentrations a unique state is observed having FRET values between those of the open and closed states. This is probably caused by the decreased lifetime of the closed, effector bound state. (**D**) Histogram of the FRET states of 𝛿IC2 at the indicated effector concentrations. The histogram contributes a Gaussian distribution with its peak gradually shifting from the peak of the open to that of the closed state. Such results are indicative of fast transitions in the sub-millisecond time scale. Lifetimes in the sub-millisecond time scales are in excellent agreement with the MD trajectories (Fig. 1E), showing effector driven 𝛿IC2 transitions to a semi-closed state (fig. S4K) with a lifetime of ~50 ns, extremely short-live to be detected by almost any experimental approach. Therefore, we conclude that 𝛿IC2 can close, however with an extremely short lifetime.

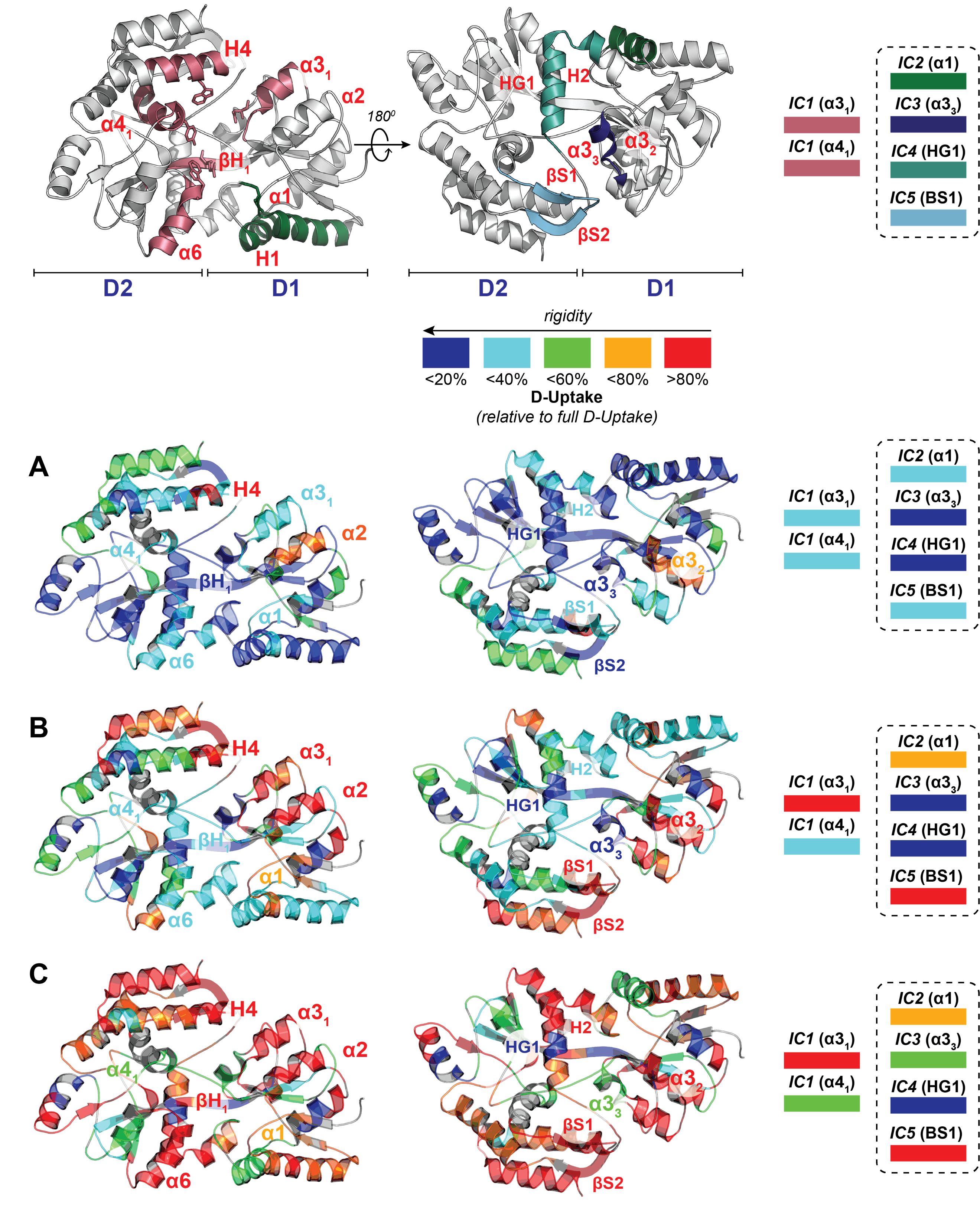

**Figure S3. HDX-MS uncovers the distribution of rigidity-flexibility of the apoprotein state.** HDX-MS enable us to probe Tier-1/2 dynamics with near residue level (~5 residues) resolution. The secondary and tertiary structure of the protein is stabilized by hydrogen bonds involving backbone amide hydrogens (NHs). Any structural fluctuation will disrupt such a hydrogen bond network, rendering the NHs solvent accessible, exchanging hydrogen with deuterium from the solvent (D_2_O). MalE was isotopically labelled (pD 7.4, T=25°C) for different periods (10, 10^3^ and 10^5^ sec). The degree of deuterium incorporation relies on the ~1 Da difference between ^1^H and D and is determined by mass spectrometry after proteolysis. The peptic fragments covered 95% of the whole protein sequence. Deuterium uptake values are indicated for the incubation times, i.e., 10 sec **(A)**, 10^3^ sec **(B)**, and 10^5^ sec **(C)**, in deuterated buffer and conveyed relative to the fully deuterated control (see also fig. S5 for plots of the relative of D-uptake *vs.* time). These values are mapped onto PDB 1OMP using the indicated color gradient. Proline residues as well as the first residue of each peptide were excluded from mapping, as they do not contribute to the observed D-uptake. (**A**) In the shortest D pulse (10 sec), the structural elements having almost complete H/D uptake include helix α2 and α3_2_. Such elements belong to D1, and are detached from the structural core, thus more accessible to the solvent. (**B**) The subsequent D pulse (10^3^ sec) uncovers the IC structural dynamics. Such correspond to the islands with different stability, as described by Ernesto Freire *et al* (*31*) to be the determinants of allosteric signal transmission. The secondary structure elements belonging to IC1,3&4 (e.g., α3_3_, α4_1_, α6 and H4), maintain an elevated degree of rigidity. Contrary, the ones belonging to the D1 (α3_1_ of IC1 and α1 of IC2) and the hairpin of IC5 are extremely flexible. (**C**) The latest D pulse (10^5^ sec) reveals the firm core of the CC. This includes IC1 (α4_1_) and IC4 (HG1). As shown in figure S1, Y155 is the key residues that stabilize effector docking. Increased rigidity is indeed required for the “Lock-and-Key” type of mechanism (*32*). The key residue Y155 (extremely conserved, can only vary to F) is responsible for placing the reducing end of maltose (effector) on the docking domain D2. On the other side, M321 is the key residue to stabilize the open state (*2*). Complete data sets are located in Auxiliary files

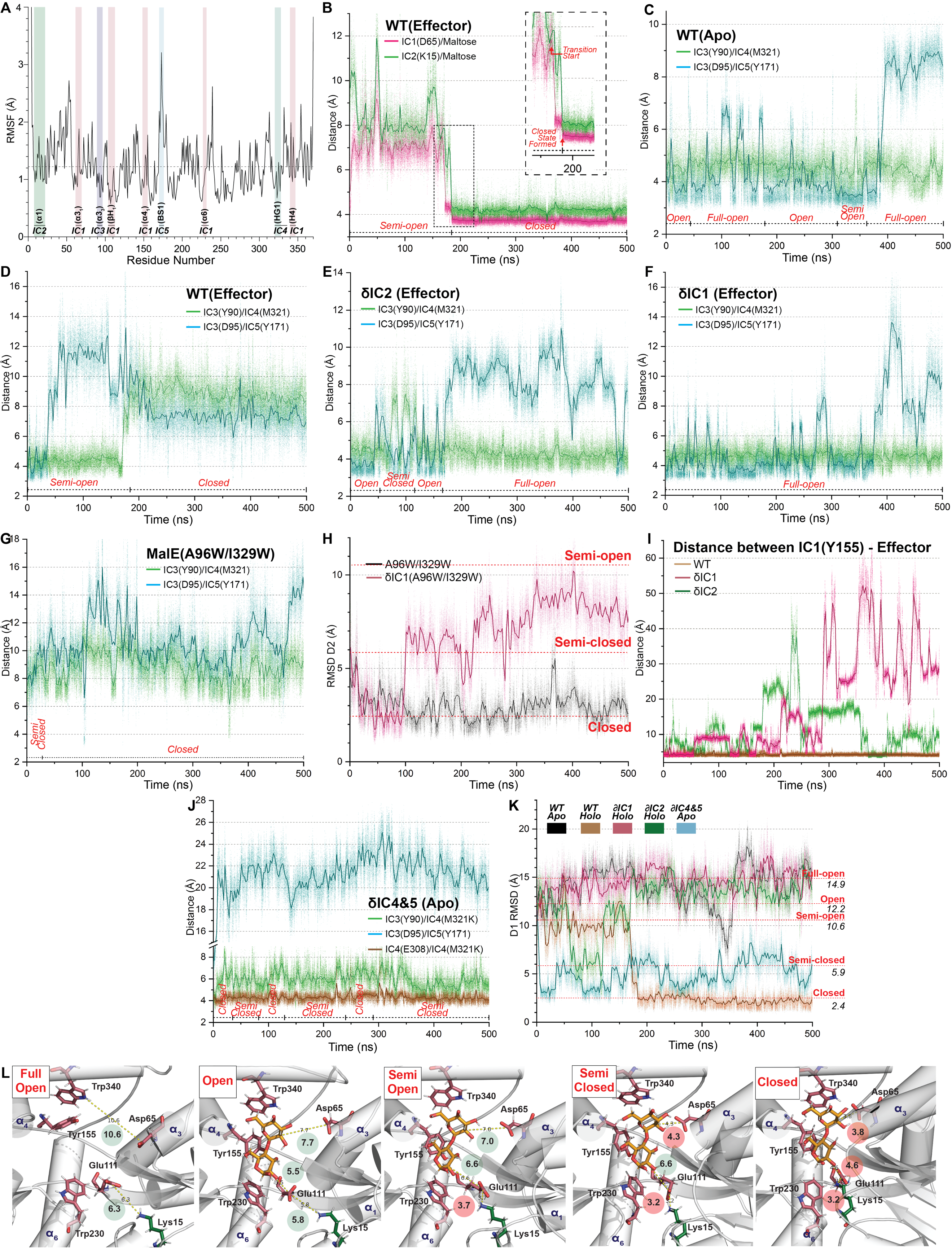

**Figure S4. MD exposes the intrinsic IC dynamics and those triggered by the allosteric effector:** To examine the interplay between Tier-1/2 and the Tier-0 state transition dynamics together with the role of the ICs in the allosteric cascade, we performed all-atom molecular dynamic simulations. We used the structural information of the open state (retrieved by extracting the protein structure from PDB: 1LLS) as the starting coordinate for our apoprotein condition. We determined the protonation state of each charged residues at pH 8 using the H++ webserver. Then, we docked on this structure maltose, being the effector (modified from PDB: 1EZ9) to produce the complex of the open effector-state. Subsequently, we set the parameter for such a complex using the FF14SB & GAFF forcefields. We carried out the MD simulations using AMBER20 under periodic boundary conditions (PBC) at 302K with an explicit water model (TIP3P) for 500ns. Finally, to evaluate the conformational transitions, we determined distance fluctuations of D1. For this, we computed the Root Mean Square Deviation (RMSD) of D1 after superimposing D2 to the crystal structure of the closed-effector state (PDB: 1ANF), used as the reference. Moreover, to determine the crosstalk between the ICs, we correlated the overall structural changes with distance changes between the ICs. In brief, the protein in the apostate undergoes subtle structural changes, in which the RMSD of D1 fluctuates from 13.7 Å to the lowest average of 10.7 Å (see also Fig. 1E – black). This result is also in line with our previous single-molecule FRET observations (*2*) and prior MD results (*9, 33*), indicating that the protein is present in an open apoprotein-state, with no observable intrinsic transitions to the closed one. (**A**) Root Mean Square Fluctuations (RMSF) of MalE residues during apoprotein-state simulation: Such a calculation provides insights regarding the molecular flexibility of GCCPs, by extrapolating the Root Mean Square Fluctuations (RMSF) of each residue. Residues belonging to IC1&4 have on average low RMSF values (i.e., below the dotted line), whereas those of IC5 increased ones, indicative of highly rigid and flexible regions respectively. IC3&2 have intermediate values denoting rigid and flexible regions respectively. Such observations are in full agreement with the HDX-MS analysis (shown in fig. S3). This indicates that MD is in line with the experimental observations with respect to the D1-D2 large-scale domain-domain motions (Tier-0 dynamics, see Fig. 1E *vs.* Fig. 2A-D), but also to the localized fluctuations (Tier-1/2 dynamics). (**B-G, J**) Distance fluctuations of the ICs by reporting distances between key representative residues together with the large-scale structural changes (Tier-0 dynamics) retrieved from panel K and reported at the bottom of each panel. (**B**) Proximal ICs-effector interactions for allosteric signal propagation: The trajectory in the presence of the effector reveals the large-domain (Tier-0) movement of D1 towards D2 (in agreement with smFRET, Fig. 2A), starting at ~170 ns. IC1 residue D65 gets in full contact with one of the glucosyl rings of the effector (magenta line, Movie S2) causing the D1-2 closing motion. The fully closed state is obtained after K15 of IC2 binds the second glucosyl ring (green line). (**C**) Distal ICs behave distinctively: In the apoprotein state simulation, fluctuations between the full-open and the semi-open states are observed. During such intrinsic dynamics, IC3 remains tightly packed to IC4, whereas IC5 undergoes cycles of attachments/detachments (see also movie S1). The rigidity of IC3&4 versus the flexibility of IC5 (fig. S4A and fig. S3) contributes to the dynamics between the ICs. (**D**) Distal IC detachment triggered by the allosteric event: Effector-IC1 interaction triggers complete detachment of the distal ICs. IC4 is fully detached from IC3 concomitantly to the effector's tight association on IC1 (panel B). On the other hand, the flexible IC5 is detached way before the allosteric (closing) event. The flexible IC5 undergoes detachment events alike at effector-free conditions (panel C, see also movie S2). (**E**) Distal ICs are temporarily detached in 𝛿IC2: Effector association on 𝛿IC2 triggers acquisition of a semi-closed state (panel K), and only during this time period, the critical IC3-4 interface is disrupted (see also movie S5**)**. (**F**) Distal ICs fail to get detached stably in 𝛿IC1: The rigid IC3-IC4 interface remains tightly packed, despite the presence of the effector. The flexible IC5 undergoes detachment events alike at effector-free conditions (panel C, see also movie S4). (**G, H**) IC1 is necessary for stabilizing the intrinsically obtained closed state: A96W/I329W destabilizes the IC3-4&5 interfaces (G) causing the closed state transition within 50 ns of simulation, alike in 𝛿IC4&5 (panel J *vs.* G). 𝛿IC1(A96W/I329W) transits to a short-live closed state (H), rationalizing the smFRET results (see fig. S2A for data and detailed explanation). **(I)** Proximal IC disruption affects the lifetime of the effector-docked state: The lifetime of the effector-docked state is a critical determinant for allosteric signal propagation (*32, 34*). To extrapolate the lifetime of this state from the MD trajectories, we calculated the distance between Y155 and the first glucosyl ring of maltose, as Y155 is the critical residue for effector docking (fig. S1, C to E and fig. S3). 𝛿IC1 or 𝛿IC2 severely compromise the lifetime of such a state, impeding stable effector docking on the cleft. As also shown by HDX-MS, due to D65A mutation in 𝛿IC1, the IC1/α6 peptide loses its rigidity (fig. S5H), compromising the stable effector docking to the cleft and driving 𝛿IC1 to the full-open state (panel K). The effector association on D2 in 𝛿IC1 is compromised starting at ~200ns of the simulation and unbinds completely from D2 at ~300 ns. (**J**) 𝛿IC4&5 bypasses the need for an allosteric trigger: 𝛿IC4&5 transits readily to the closed state, irrespective of the presence of the effector (see also Movie S3**)**. The critical to the allosteric event IC3-4 interface is intrinsically disrupted. M321K (IC4) perturbs the IC3-4 hydrophobic patch and interacts electrostatically with the IC4 residue E308. (**K, L**) Critical interactions on the effector-cleft stabilize the five states of GCCP: Combined MD results of apoprotein- and effector-states gave us detailed insights into how the cleft interactions stabilize the five MalE states. The average RMSD of the five states is also indicated. The number of interactions between IC1&2 residues and the effector correlates with the formation of five different states, i.e., full-open (FO), open (O), semi-open (SO), semi-closed (SC), and closed (C). C, SC, SO, and O/FO states are stabilized by three, two, one, and zero interactions, respectively. **(L)** As fluctuations from C to SC and O to SO alter marginally the distance between the two domains (table. S2) and occurs at ns time-scales, such are challenging to be probed by smFRET. The interacting residues are indicated by red circles followed by a number representing the distance in Angstrom (Å).

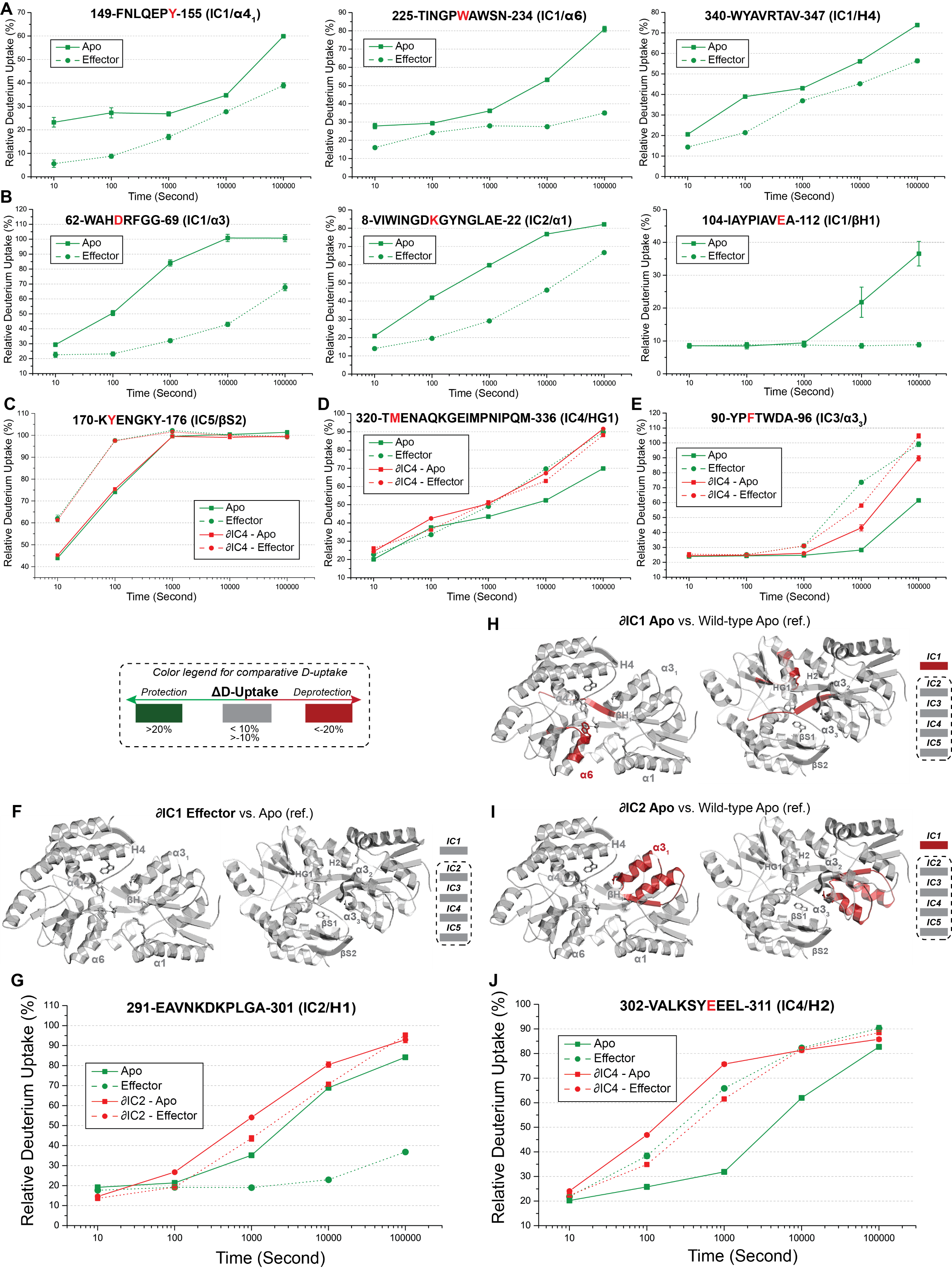

**Figure S5. HDX-MS reveals the localized (Tier-1/2) dynamics upon effector association.** To identify the overall dynamics triggered by the allosteric event, we performed HDX-MS analysis at different conditions, as indicated. (**A, B**) Relative deuterium uptake for the IC1&2 cleft peptides of D2 (A) and D1 (B) as a function of deuterium incubation. Upon effector association, the D-uptake is reduced in the binding cleft. This could be attributed to direct interaction with the effector, reduced solvent accessibility, re-enforcement of pre-existing contacts or stabilization by new contacts in the closed state. Among the effector binding residues, only Y155 makes a backbone hydrogen bond with the effector (fig. S1) in line with the significant protection of peptide 149-155 (panel A, left) at the earliest time point. (**C-E**) Relative deuterium uptake for the distal-IC peptides over time upon effector binding: Effector association increases dramatically the dynamics of the IC3,4&5 peptides. (**C**) As detailed in the main text, we observed an elevated degree of IC5 dynamics induced by the effector (see also Fig. 2E**)**: the IC5 hairpin uptakes deuterium an order of magnitude faster compared to the apo-state. These observations agree with the MD results (see movie S1 *vs.* movie S2 and fig. S4D) and in line with the role of the hairpin for stabilizing the open apo form of the thermophilic MalE of *Thermotoga maritima*, decrypted by MD simulations (*35*). (**D**) The ‘rigid IC4’ (see also fig. S3) undergoes a significant structural loosening triggered by the effector, represented in the exposure of the HG1 peptide to deuterium. (**E**) The increase in deuterium uptake of the two distal ICs is observed in IC3, i.e., the α3_3_ peptides. The disruption of the critical IC3-4 interface in 𝛿IC4 destabilizes the IC3&4 peptides (i.e., increases their flexibility) similarly to the effector (panels D&E). Contrary, it has no influence on the IC5 peptide (panel C). Thus, we did not observe any cross-talk between the distal ICs (4&5). Both interfaces (i.e., IC3-IC4& IC3-IC5) need to be destabilized (𝛿IC4&5) for allosteric signal propagation, by-passing the need of an effector (Fig. 2B, Fig. 1E, see also main text for details). We conclude that effector association on IC1&2 destabilizes allosterically the IC3-IC4&5 interfaces. (**F**) IC1 is necessary to trigger allosterically the destabilization of the distal ICs: Comparative deuterium uptake analysis of the 𝛿IC1-effector complex compared to the apo-state. Upon saturating effector concentrations (i.e. 50 mM maltose; fig. S6A), 𝛿IC1 does not undergo statistically significant structural changes in all identified peptides. (**G**) IC2 rigidification is essential for the acquisition of a stable effector-driven closed state: Because of the K15A mutation, it is impossible to compare the absolute deuterium uptake of the α1 peptide (where K15 resides) upon effector binding. To probe the allosteric event on IC2, we compared deuterium uptake of H1. H1 is located near α1 and both helices move simultaneously with D1 upon closing. H1 of IC2 is rigidified upon effector association (green lines) alike α1 (see panel B-middle). 𝛿IC2 fails to transmit allosterically this effector-driven rigidification (red lines). As 𝛿IC2 is proficient in transmitting allosterically destabilization of the IC3-4&5 interface (Fig. 2F and fig. S4E), we conclude that H1 rigidification modulates the lifetime of the closed states (see also fig. S2C). 𝛿IC2 is capable of acquiring the short-lived semi-closed state (fig. S4K). (**H**) 𝛿IC1 reveals the intra-IC1 allosteric cross-talk: Comparative deuterium uptake analysis of the apoprotein-state of 𝛿IC1 compared to the wild-type. Disruption of IC1 on D1 (i.e., D65A) abrogates the rigidity of IC1 cleft residues (on α6 and βH1) that are critical for effector docking (fig. S1, C to E). Thus, the effector is not stably docked on the cleft in 𝛿IC1, in line with the MD analysis (fig. S4I). (**I**) 𝛿IC2 reveals the inter-IC1/IC2 allosteric cross-talk: Comparative deuterium uptake analysis of the apo-state of 𝛿IC2 compared to the wild-type. Disruption of IC2 eradicates the rigidity of key IC1 residues (fig. S3, B and C and fig. S4A) for effector docking (fig. S1, C to E). Once more, this explains the unstable effector docking on 𝛿IC2 (fig. S4I). (**J**) The destabilization of the distal ICs upon effector association is reversed in 𝛿IC4: IC4 disruption (red straight line) renders H2 of IC4 intrinsically destabilized, alike upon effector association on wild-type (green dashed line), similarly to HG1 of IC4 (panel D). However, H2 is rigidified upon effector association on 𝛿IC4 (red dashed line). Such a rigidification stems from the electrostatic interaction between the K321 (in 𝛿IC4) and an H2 residue (E308), as shown by MD (fig. S4J and movie S3). Complete data sets are located in Auxiliary files.

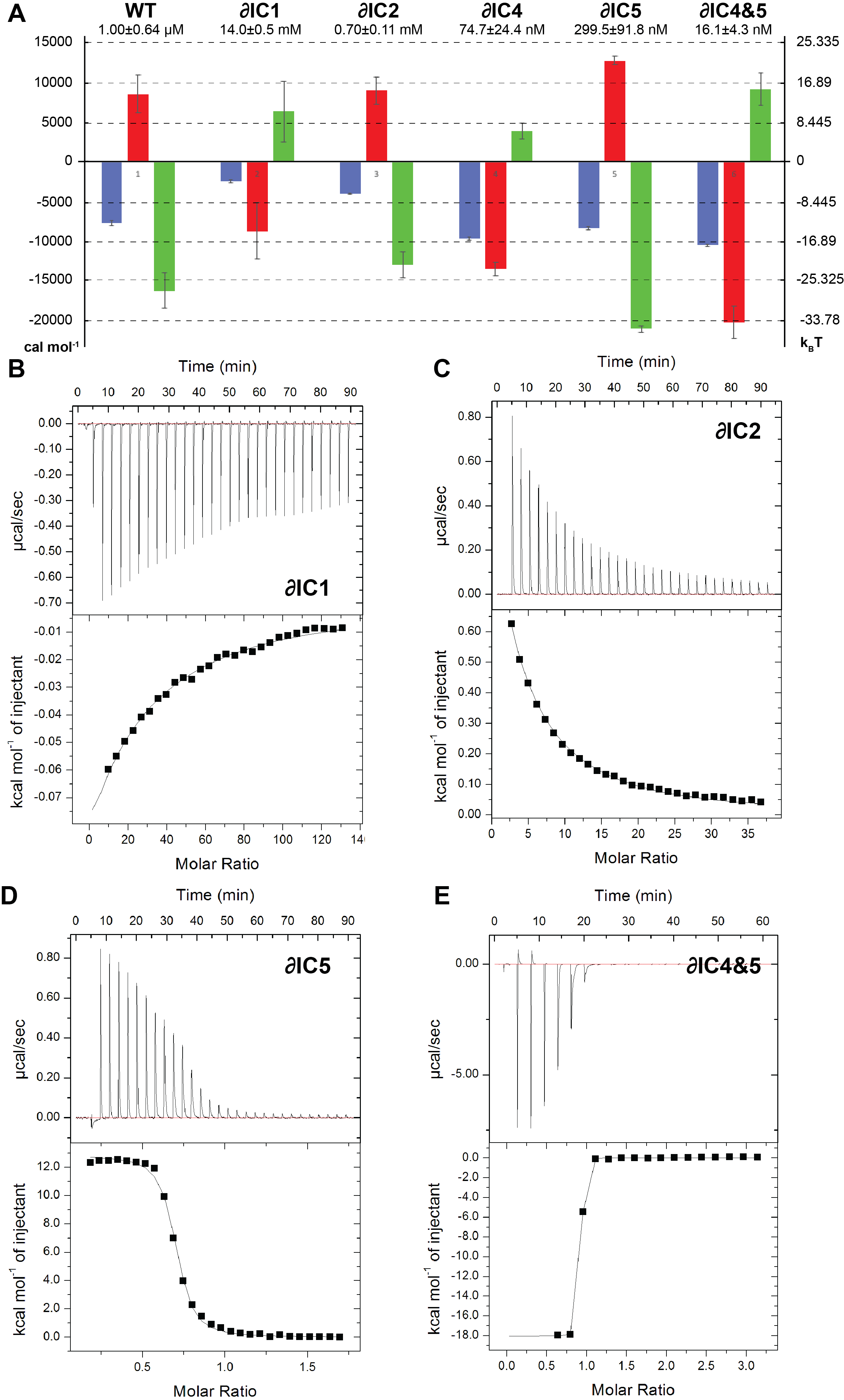

**Figure S6. Thermodynamics of effector association determined by ITC.** (**A**) Summary of the thermodynamic parameters of effector association to the indicated derivatives. The calculated binding affinities of the derivatives are depicted (n=3) together with the standard error of the mean (SEM). An indicative thermogram of 𝛿IC1, 𝛿IC2, 𝛿IC5, and 𝛿IC4&5, upon effector binding is shown (**Β-Ε**). Experiments were carried out at 7^0^C (or 25^0^C for 𝛿IC4 and 𝛿IC4&5). G, ΔH, and -TΔS are indicated respectively with blue, red, or green bars. 𝛿IC1 causes a four-order reduction in effector affinity. Importantly, 𝛿IC1 exhibits the thermodynamics signature (i.e., enthalpically driven binding) of proteins utilizing aromatic residues to bind carbohydrates (*36, 37*), exploiting the relatively weak CH/π interactions. As 𝛿IC1 fails to trigger the structural transition to the closed state (Fig. 2C), the reduced affinity originates from the weak “docking” interaction of maltose exclusively with the D2 IC1 residues (fig. S1C).

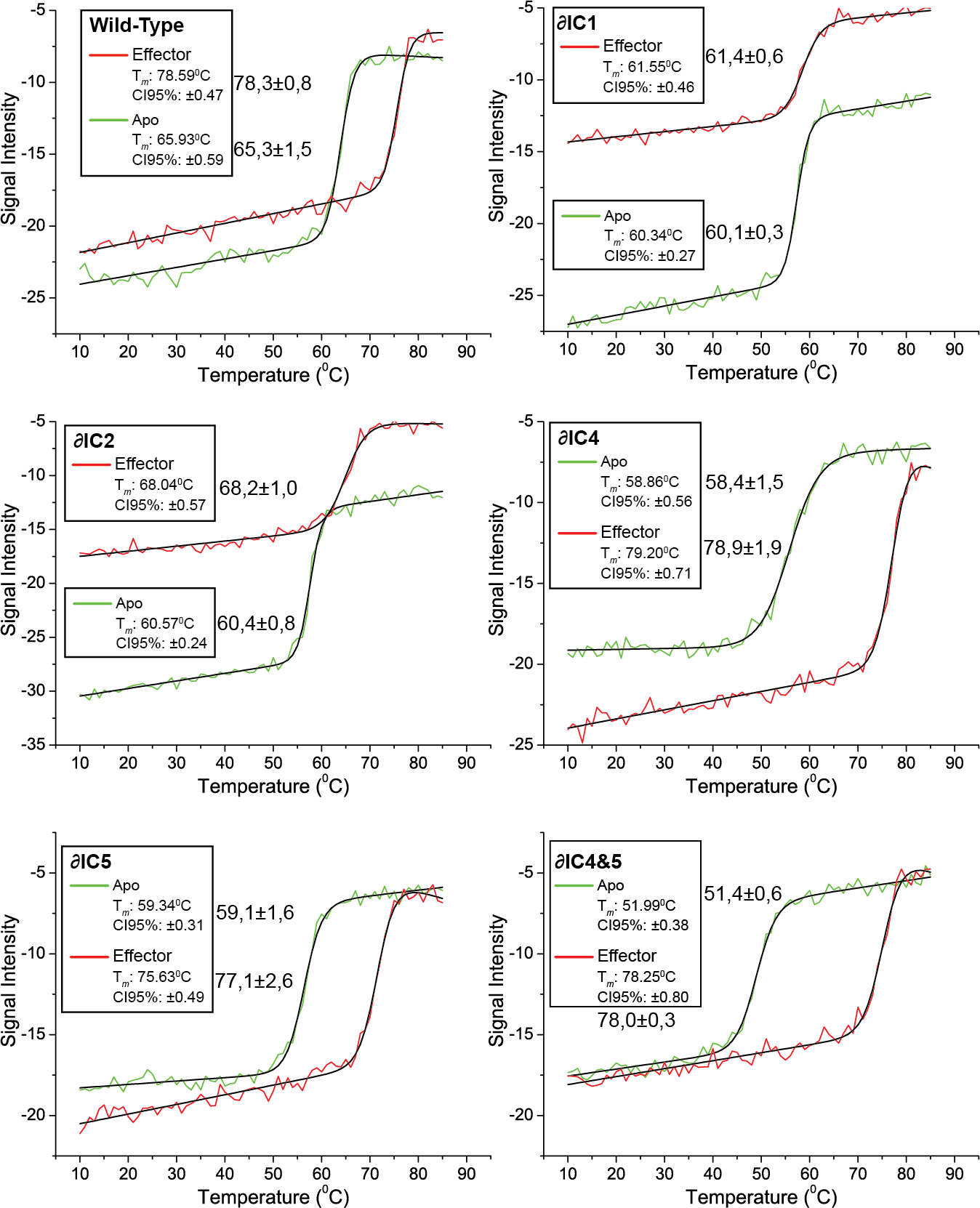

**Figure S7. CD reveals the influence of IC disruptions and effector association on the thermodynamic stability.** The indicated derivatives have been subjected to thermal unfolding experiments at a temperature range of 10 to 90^0^C in the presence or absence of the effector (green and red lines, respectively). The signal was monitored at 280 nm (for the details see Materials and Methods). The fitting (black lines), melting temperatures (T*_m_*) and the 95% confidence interval (CI95%) have been retrieved from the CalFitter webserver (*15*) after fitting the experimental data with default parameters (20 times iteration) using N=D (Van’t Hoffs) model. Melting temperatures together with the standard error of the mean (SEM) are also indicated (n=3).

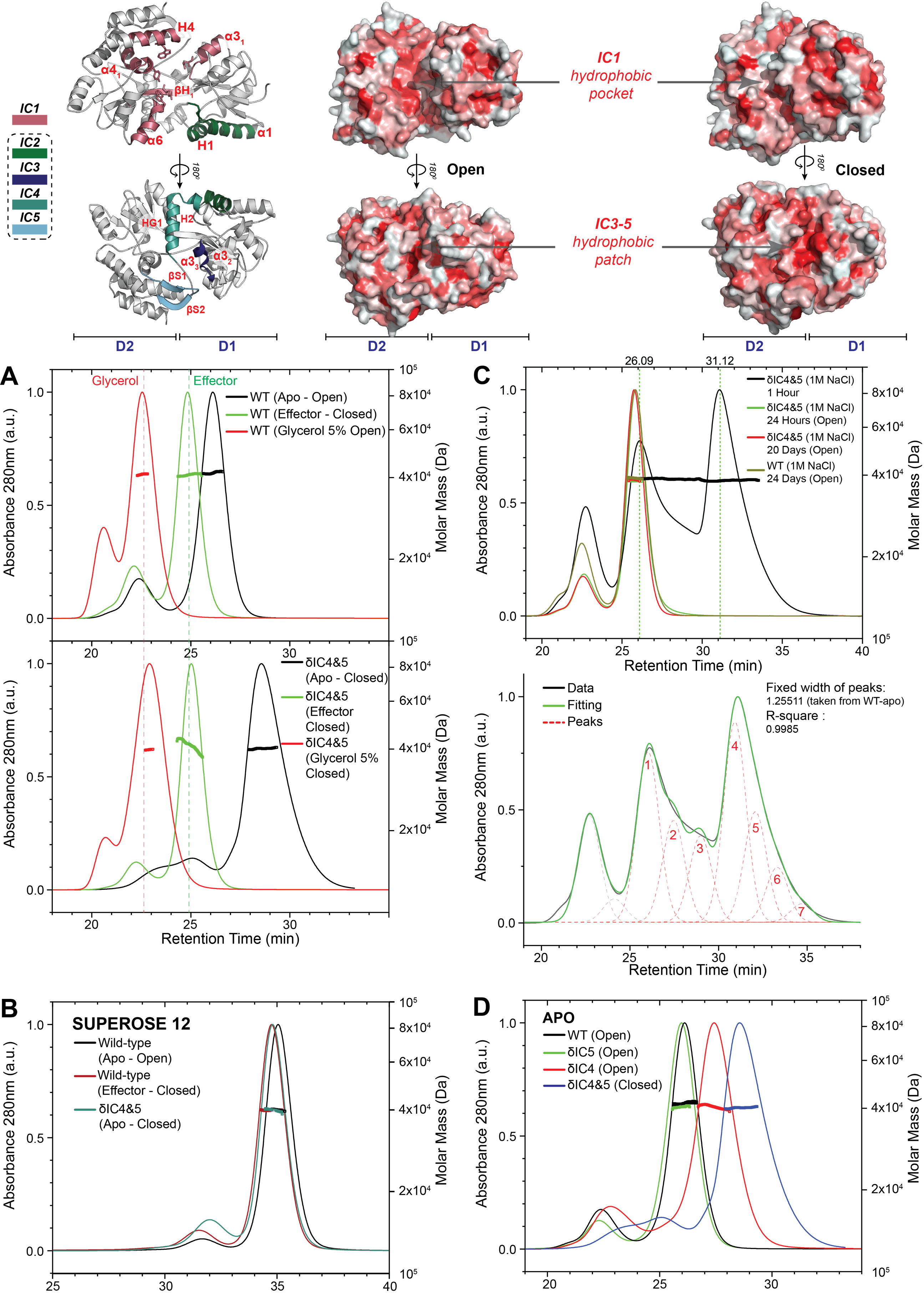

**Figure S8. Distinct structural states are physically separated based on their hydrophobicity. *Physical separation and effector size:*** To physically separate the intrinsically obtained structural states and those induced by effector-association, we analyzed meticulously the (structural) properties of them. Effector docking occurs on IC1 residues belonging to D2, and specifically on the aromatic residues (Y155, W230, W340) that stabilize effector association to the open state (fig. S1, C to E). The aromatic residues belong on elements having increased rigidity (fig. S3C); hence, this initial encounter with the effector follows the properties of a typical “lock-and-key” mechanism (*34*). In such a case, the interaction of a ligand with a rigid domain maximizes the entropic gain ΔS_solvation_, known as the hydrophobic effect and analogous to the first folding step to obtain the molten-globule state. Clearly, the amount of water molecules displaced (strength of the hydrophobic effect) due to the protein-effector surface complementarity correlates directly with the size of the effector. Thus, the first energetic obstacle -by-passed by the hydrophobic effect- to reach the transition state (Fig. 3A) correlates with the size of the effector (see fig. S9 and main text for details). This activation barrier depends on the energetic level of the open state, modulated during evolution primarily by the distal ICs3-5 (Fig. 3A). IC3-5 in the open state are tightly packed alike a hydrophobic patch (fig. S1F). In the effector driven closed state or in δIC4&5, the hydrophobic patch is ruptured and exposed to the solvent (Fig. 2, E and I and fig. S4, D and J). This enthalpy penalty is compensated by the increased entropy gain due to ΔS_configurational_ (fig. S6A) derived from the increased structural dynamics in IC3-5 (Fig. 2E and fig. S5, C to E). Thus, effector association hinders the IC1 hydrophobic pocket, whereas the IC3-5 hydrophobic patch gets exposed in δIC4&5, as also previously documented (*38*). We opted to rely on these properties to physically separate structural states, by size-exclusion chromatography (SEC) on Sepharose 75, a resin-material known to bind slightly hydrophobic patches (*39, 40*). We indeed succeeded to separate proteins based on their hydrophobicity and not relying on their hydrodynamic radius, as we verified the protein size by multi-angle light scattering (MALS). For this we here present the most indicative results: **(A**) i. Chromatogram of wild-type (upper panel) and 𝛿IC4&5 (lower panel) at the indicated conditions, in the presence or absence of the effector. The effector hinders the IC1 hydrophobic pocket, and the effector-bound protein elutes earlier that the retarded apoprotein state (green *vs.* black line in A, upper panel). Clearly, the two species separate because of their -retarded- elution caused by hydrophobicity and not based on hydrodynamic radii. That is why the effector-driven closed state elutes earlier than the open one, despite its reduced hydrodynamic radius. ii. 𝛿IC4&5 having exposed the IC3-5 hydrophobics (as shown by MD in figure S4J and HDX-MS in Figure 2I) manifests a retarded elution (black line in panel A**,** lower panel *vs.* black line in A, upper panel). As in the effector-driven closed state, the IC3-5 hydrophobic patch of 𝛿IC4&5 is similarly exposed to that of wild-type; the two effector-bound states elute at similar time points (green line in A**,** lower panel *vs* green line in A, upper panel). The presence of glycerol to nullify the Sepharose 75 hydrophobic interactions abolishes retarded elution and all structural states elute at the same point according to their hydrodynamic radii (red line in A**,** lower panel *vs* red line in A, upper panel). (**B**) The separation relying on retarder elution due to non-specific hydrophobic interactions with Sepharose 75 are completely lost in other resin materials, i.e., Superose 12. **(C)** The free energy landscape (FEL, e.g., the one presented in Fig. 3A) is modulated by thermodynamic (e.g. denaturants, temperature, pH, ionic strength) or functional (e.g. effectors, partners, perturbations) properties, bringing out the non-complementary entropy-enthalpy change within protein-solvent systems (*41*). Actually, the free energy wells and energy barriers depict the static forms of manifestation of such changes (*42*). As acquisition of distinct structural states (i.e., energy wells) are governed by the exposure of hydrophobic patches (panels A, B), we analyzed further the apoprotein state of 𝛿IC4&5, manifesting the greatest retardation in its elution. This is caused by the fact that in 𝛿IC4&5 at apoprotein conditions, the IC1 and IC3-5 hydrophobic patches are fully exposed. To perturb drastically the FEL of apo 𝛿IC4&5, we increased the ionic strength (1M NaCl) and performed SEC-MALS. Due to this drastic change in the protein-solvent system, the entropy-enthalpy equilibrium was reached in 24 hours (green and red lines, panel C). The retarded apoprotein 𝛿IC4&5, present in the closed state at physiological salt concentrations (panel A**)** equilibrates to the apoprotein open state after 24 hours. Since the kinetics to the “new” equilibrium are very slow, we managed to “freeze” several distinct (>5 judging from the amplitude of the chromatogram) structural states (black line, lower panel in C). Similarly, a number of states also emerged in the MD trajectories from the open to the closed state (fig. S4, K and L); showcasing that the FEL can be modulated by different factors, i.e., effectors/perturbation. (**D**) Chromatogram of wild-type and the 𝛿IC4 or 𝛿IC4&5 derivatives at apoprotein conditions.

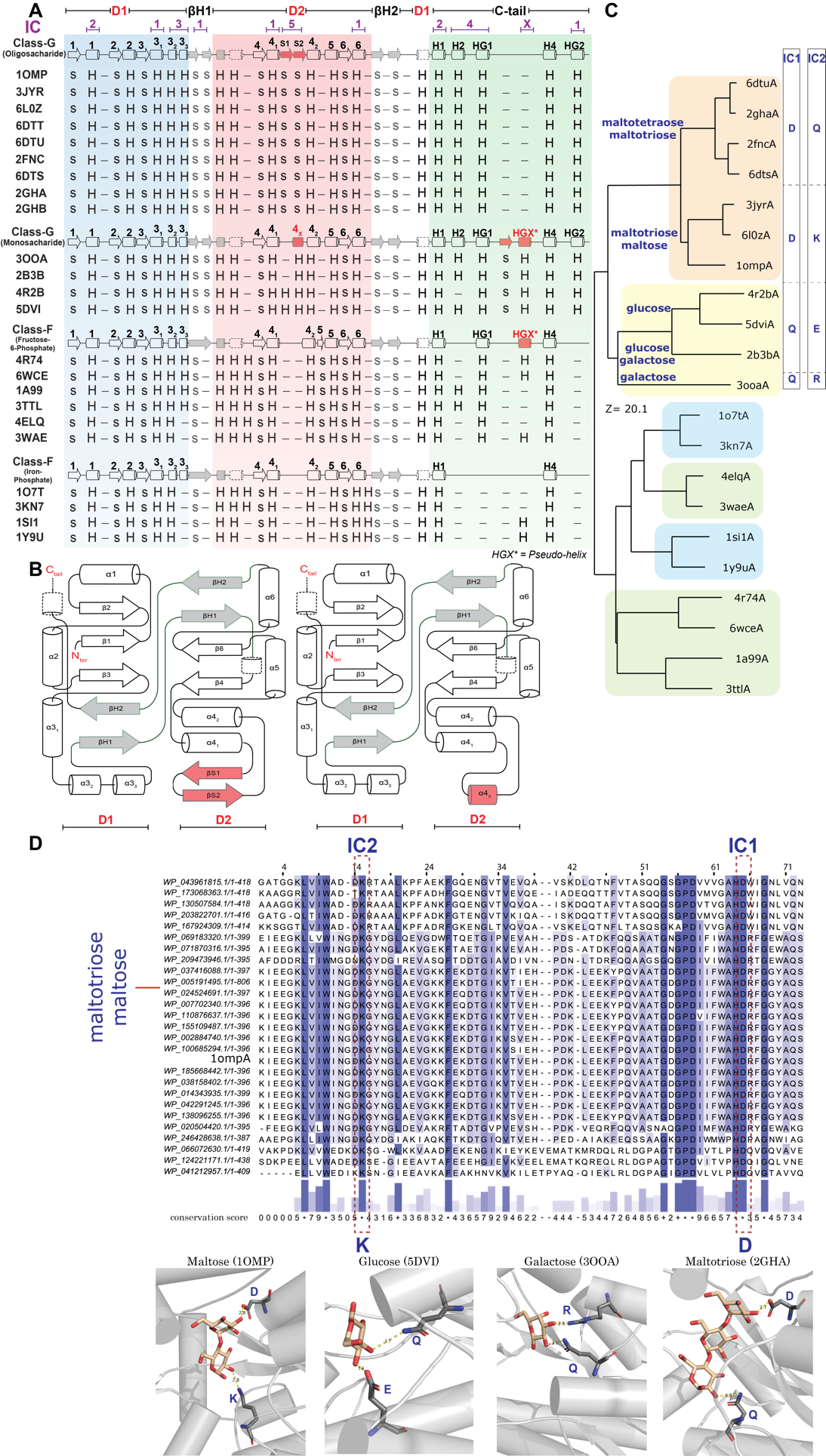

**Figure S9. The IC modules dictate effector specificity and environmental adaptation.** (**A**) Secondary structure (SS) alignment of various CCPs, including oligosaccharide & monosaccharide GCCPs and the fructose/glucose-6-phosphate, putrescine/spermidine and thermophilic class-F CCPs (FCCPs). Secondary structure elements marked in red represent modules differing in CCPs recognizing structurally dissimilar effectors. When monosaccharides act as the effectors, GCCPs have not retained IC5 (βS1&2) while acquiring the pseudo-helix HGX. Interestingly, a subcategory of the class-F CCPs that accepts large effectors as fructose/glucose-phosphate, i.e., AfuA&P5PA (PDB: 4R74&6WCE); putrescine/spermidine, i.e., PotF&SpuE, (PDB: 1A99&3TTL) or belong to thermophilic organisms i.e., Thermus thermophilus (PDB: 4ELQ, 3WAE) embrace the IC4 module (HG1). As AfuA&P5PA bind monosaccharides, they possess the HGX module. As previously published, FCCPs are missing the IC5 modules **(*2*)**. (**B**) SS topology of D1 and D2 highlighting the missing IC5 modules between GCCPs that bind oligosaccharides (left) or monosaccharides (right). (**C**) Structure-based phylogenetic tree with indicative class F & G structures. The simplified phylogenetic tree is shown in Figure 3C. Conserved IC1 and IC2 residues of each structural category (i.e. distinct branches) are indicated in the right side of the panel. (**D**) A representative sequence alignment of the MalE (1OMP) structural category showcasing the conserved IC1 and IC2 residues (upper part). Such conserved residues vary depending on the effector, as evident on panel C. In the bottom part, a representative structure in complex with the different effectors indicates that the IC1 and IC2 key residues evolved to adapt for effector binding.

Table S1. Mutations disrupting the ICs used in this study.

| **Mutant ID** | **Mutations** | **Reference** |
| --- | --- | --- |
| 𝛿IC1 | D65A | Introduced here |
| 𝛿IC2 | K15A | Introduced here |
| 𝛿IC4 | M321K | Gouridis, G., Muthahari, Y., *et al*. (*2*) |
| 𝛿IC5 | Δ172, 173, 175 and 176 | Shilton, *et al*. (*43*) |
| 𝛿IC4&5 | M321K, Δ172, 173, 175 and 176 | Introduced here |

Table S2. T36-S352 distance in the 5 states revealed by MD

| **State** | **T36-S352 Distance (nm)** |
| --- | --- |
| Full open | 5.32 |
| Open | 5.17 |
| Semi Open | 4.78 |
| Semi Closed | 4.65 |
| Closed | 4.04 |

The distance between S36 and T352 in each state of GCCPs. The distance is measured from the representative structures upon KMeans clustering (see Materials and Methods).
